# Inside insight: decoding how insight emerges from competing world models

**DOI:** 10.64898/2026.05.21.726889

**Authors:** Kengo Inutsuka, Tadaaki Nishioka, Tom Macpherson, Mana Fujiwara, Takatoshi Hikida, Honda Naoki

## Abstract

When and how does insight emerge? We conceptualize insight as a sudden realization arising from restructuring a world model: an internal interpretation linking actions to outcomes. Yet these latent dynamics remain difficult to access, even with behavior and verbal report. Here we developed inside insight dynamics (IID), a machine-learning framework that estimates latent world-model dynamics from behavioral data. Using IID, we analyzed mouse behavior in indirect- and direct-rule tasks, each requiring a shift from an initial world model to a rule-consistent representation. IID inferred the “when” of insight-like shifts by estimating the timing of transitions between competing world models, and examined the “how” by comparing alternative learning processes underlying them. This analysis revealed distinct mechanisms of world-model learning: the indirect- and direct-rule tasks were better explained by gated learning and parallel learning, respectively. Thus, IID opens a route to quantifying latent insight dynamics from observable behavior alone.

## Introduction

In daily life, flexible problem solving often requires more than the gradual refinement of an existing strategy. In some situations, a solution emerges only after a qualitative change in how the problem itself is represented [1, 2]. Such insight-like change is often described as an “Aha!” moment, in which an individual abruptly shifts from one interpretation of a situation to another [3]. A useful historical analogy is a conceptual shift from a geocentric to a heliocentric view of planetary motion: the same observations become explainable once the underlying representational framework changes [4]. Although insight has long been studied in psychology and cognitive neuroscience, its underlying neural and computational mechanisms remain difficult to define quantitatively [2, 3].

One way to formalize insight-like change is as a restructuring of an internal “world model.” A world model is an internal representation of how the external environment is organized and how actions lead to outcomes [5, 6]. From the perspective of the Bayesian brain hypothesis, such a model can be viewed as a generative model that enables an agent to infer hidden causes of observations and predict future outcomes [7, 8]. In this framework, insight-like change can be understood as a transition between competing world models: the agent stops interpreting the environment through one model and begins to rely on another model that better explains the observed action-outcome relationship.

However, this view raises an important computational question. If multiple candidate world models exist, how are they learned and selected [6, 9]? One possibility is a gated learning mechanism, in which each world model is updated in proportion to the agent’s current reliance on that model [10]. Under this mechanism, insight-like change is not merely a sudden behavioral readout, but a process in which an emerging world model gradually becomes available for learning as it gains internal weight. An alternative possibility is a parallel learning mechanism, in which multiple candidate world models are updated simultaneously, regardless of their current contribution to behavior. In this case, an apparent moment of insight may reflect the later selection of a model that has already been learned implicitly [5]. Distinguishing between these alternatives is essential for understanding whether insight-like behavior arises from belief-dependent selective learning or from parallel latent learning followed by behavioral expression.

Insight has been extensively studied in humans, often by combining behavioral performance, brain activity, and subjective reports of the moment of insight [2, 3, 11]. However, subjective reports provide only limited access to the underlying temporal dynamics, and non-invasive measurements such as fMRI and EEG, by themselves, are not sufficient to establish circuit-level mechanisms. Animal models offer the possibility of invasive neural recording, manipulation, and genetic intervention, but they lack verbal report. This makes it difficult to identify when an animal undergoes an insight-like change in internal representation. Therefore, a method is needed to infer latent insight dynamics directly from observable behavior, without relying on subjective report.

We previously developed a visual discrimination-based learning task in mice to study cue-guided inhibition behavior, and showed that dopamine D2 receptor activity contributes to avoidance of a cue associated with reward omission [12].

In the present study, we reanalyzed two related touchscreen-based tasks from that study, which we refer to here as an indirect-rule task and a direct-rule task, corresponding to the original VD-Inhibit and VD-Attend tasks, respectively (**Fig. 1a**). In the indirect-rule task, one fixed cue was consistently associated with reward omission, whereas other randomly presented cues were rewarded; therefore, mice had to learn to inhibit responses to the fixed cue and choose the alternative cue. In the direct-rule task, by contrast, one fixed cue was consistently rewarded, whereas the other randomly presented cues were unrewarded; therefore, mice had to learn to choose the fixed cue. Because cue positions varied across trials in both tasks, successful performance required learning a cue-based reward rule rather than a simple left-right choice rule. Thus, the two tasks differed in task structure: the indirect-rule task required animals to infer the rewarded choice by avoiding a specific non-rewarded cue, whereas the direct-rule task involved a more direct cue-reward association.

In this study, we developed inside insight dynamics (IID), a computational framework for estimating latent insight dynamics from trial-by-trial action and reward sequences. IID assumes that behavior is generated by competition between two candidate world models: a side-dependent model, in which reward is interpreted as depending on the chosen side, and a cue-dependent model, in which reward is interpreted as depending on cue identity. The method estimates both reward beliefs within each model and a time-varying insight variable representing their relative dominance. By comparing gated learning and parallel learning models, IID further tests how alternative world models are learned during insight-like transitions. Applied to mouse behavior, IID revealed task-dependent mechanisms: indirect-rule task was better explained by gated learning, whereas direct-rule task was more consistent with parallel learning. These results suggest that insight-like behavioral change is not governed by a single fixed computational mechanism, but depends on task structure. More broadly, IID provides a quantitative approach for inferring hidden restructuring of internal world models from behavior alone.

## Results

### Workflow for decoding insight dynamics from behavior

We first outline the overall workflow of this study (**Fig. 1**). We analyzed trial-by-trial behavioral data from two touchscreen-based visual discrimination tasks: an indirect-rule task and a direct-rule task, corresponding to the original VD-Inhibit and VD-Attend tasks, respectively. In both tasks, mice selected either the left or right option and received reward or no reward depending on a cue-based task rule (**Fig. 1a**). To explain these behavioral sequences, we constructed an agent decision-making model in which the agent internally maintains two competing world models of the task structure (**Fig. 1b**). These consisted of a side-dependent model, in which reward is interpreted as depending on the chosen side, and a cue-dependent model, in which reward is interpreted as depending on cue identity. In this decision-making model, the agent sequentially updates belief states within each world model and selects actions according to their relative dominance.

**Fig. 1.**
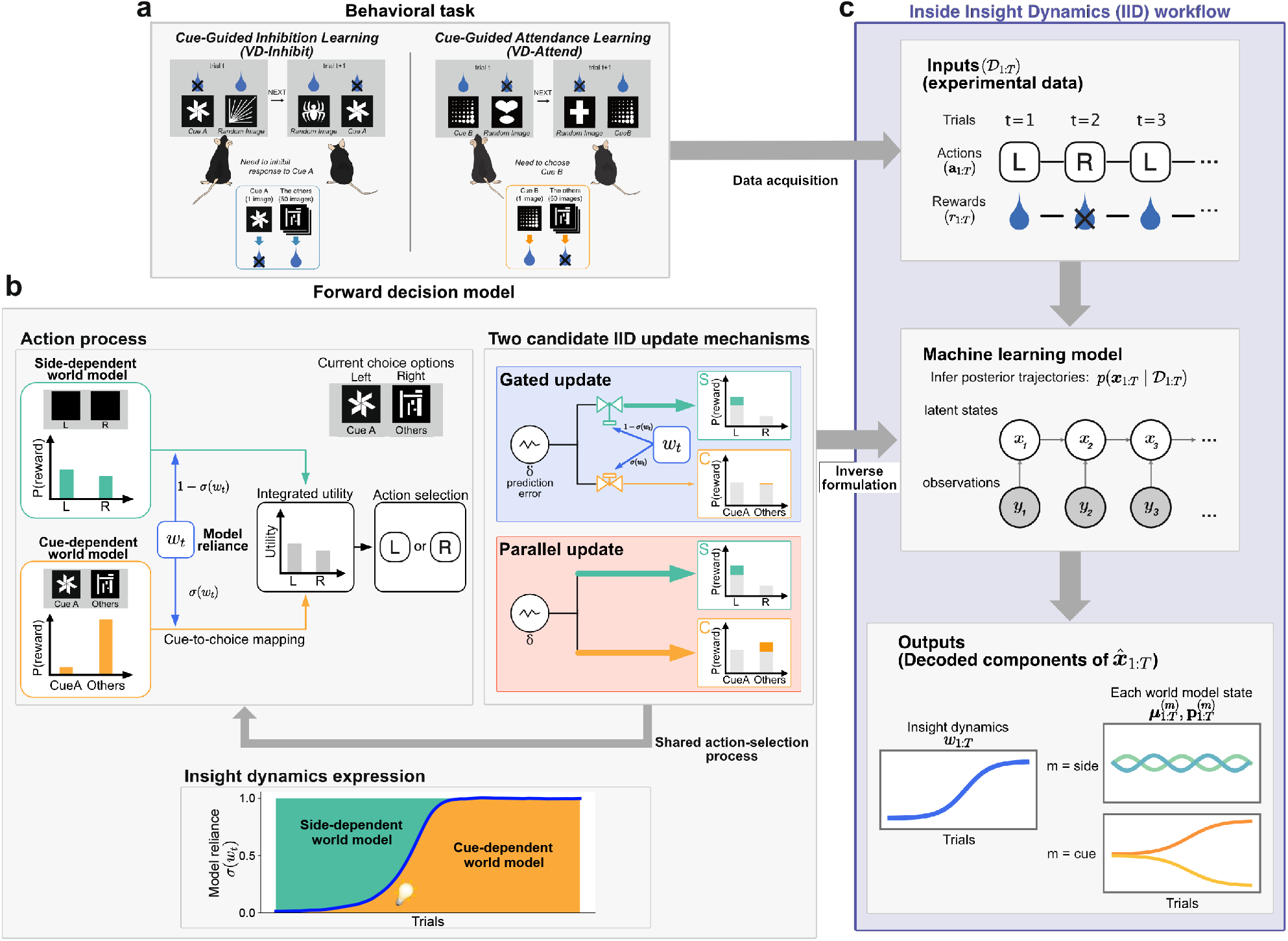
Workflow for decoding insight dynamics from behavioral time-series. **a**, Behavioral task paradigm. Mice performed two touchscreen-based visual discrimination tasks, indirect-rule task, originally termed VD-Inhibit, and direct-rule task, originally termed VD-Attend. In indirect-rule task, one fixed cue, Cue A, was consistently associated with non-reward, whereas the other simultaneously presented cues were rewarded; thus, mice had to inhibit responses to Cue A and choose the alternative cue. In direct-rule task, one fixed cue, Cue B, was consistently rewarded, whereas the other cues were not; thus, mice had to choose the matching cue. Trial-by-trial behavioral data consisted of the chosen action, left or right, and the resulting reward outcome. **b**, Agent decision-making model and candidate update mechanisms. The agent was assumed to maintain two internal world models: a side-dependent world model, in which reward is interpreted as depending on the chosen side, and a cue-dependent world model, in which reward is interpreted as depending on cue identity. Within each world model, the agent sequentially updates belief/recognition states about the reward probability for the side-dependent model and for the cue-dependent model. The agent selects actions by integrating these two world models according to a latent weighting variable *w*_*t*_, which represents their relative dominance. Insight dynamics were defined as the temporal evolution of this weighting variable, corresponding to a transition from reliance on the side-dependent world model to reliance on the cue-dependent world model. We compared two candidate update mechanisms. In the gated learning model, belief updating within each world model was weighted by the current reliance on that model. In the parallel learning model, both world models were updated in parallel, independently of their current contribution to behavior. **c**, IID workflow. The inputs to inside insight dynamics (IID) were trial-by-trial behavioral data, consisting of choices and reward outcomes, together with task-defined category mappings used to assign the cue-dependent components. IID formulates an inverse problem based on the forward decision-making model shown in **b** and estimates the latent internal trajectories that most likely generated the observed behavior. The outputs are decoded insight dynamics and decoded belief/recognition trajectories for the side-dependent and cue-dependent world models.

We defined insight dynamics as the temporal change in this relative dominance, represented by a latent weighting variable *w*_*t*_ (**Fig. 1b**). A low value of *w*_*t*_ corresponds to stronger reliance on the side-dependent model, whereas a high value of *w*_*t*_ corresponds to stronger reliance on the cue-dependent model. This relative dominance was used to define two candidate learning mechanisms (**Fig. 1b, right**). In the gated learning formulation, belief updating within each world model is weighted by the current reliance on that model. In the parallel learning formulation, both world models are updated independently of their current contribution to behavior. Thus, a transition in *w*_*t*_ represents an insight-like restructuring from a side-based interpretation of the task to a cue-based interpretation, while comparison between the two learning formulations tests how this restructuring is acquired. We then developed inside insight dynamics (IID), an observer-side inference framework that formulates the inverse problem of this decision-making model and decodes latent recognition states and insight dynamics from the observed action–reward sequences and task-defined cue-category mappings (**Fig. 1c** and **Supplementary Fig. S1**). This workflow allowed us to ask when mice shifted between world models and whether such shifts were better explained by gated learning or parallel learning.

### Behavioral transitions reflect restructuring of the world model

We next characterized the overt behavioral transitions observed during learning. In the indirect-rule task, mice initially showed an imbalance in left versus right choice probabilities, indicating that early behavior was biased toward spatial choice rather than cue identity (**Fig. 2a**). As training progressed, this side bias gradually disappeared, suggesting that the animals no longer relied primarily on a left– right choice strategy. Consistent with this transition, the fraction of correct choices initially remained near chance level but later increased to above 80% (**Fig. 2b**). The cumulative reward curve also showed a marked increase in slope, indicating an abrupt improvement in reward acquisition (**Fig. 2c**). A similar behavioral transition was observed in the direct-rule task. Mice again showed an initial imbalance in left versus right choice probabilities, suggesting that early choices were influenced by spatial bias (**Fig. 2d**). This bias was subsequently reduced as learning proceeded. The fraction of correct choices increased from near chance level to above 80% (**Fig. 2e**), and the cumulative reward curve showed a corresponding increase in slope (**Fig. 2f**). Thus, in both tasks, mice shifted from an early spatially biased strategy to a later cue-rule-consistent strategy. Individual behavioral trajectories for all mice are shown for the indirect- and direct-rule tasks in **Supplementary Fig. S2 and Fig. S3**, respectively.

**Fig. 2.**
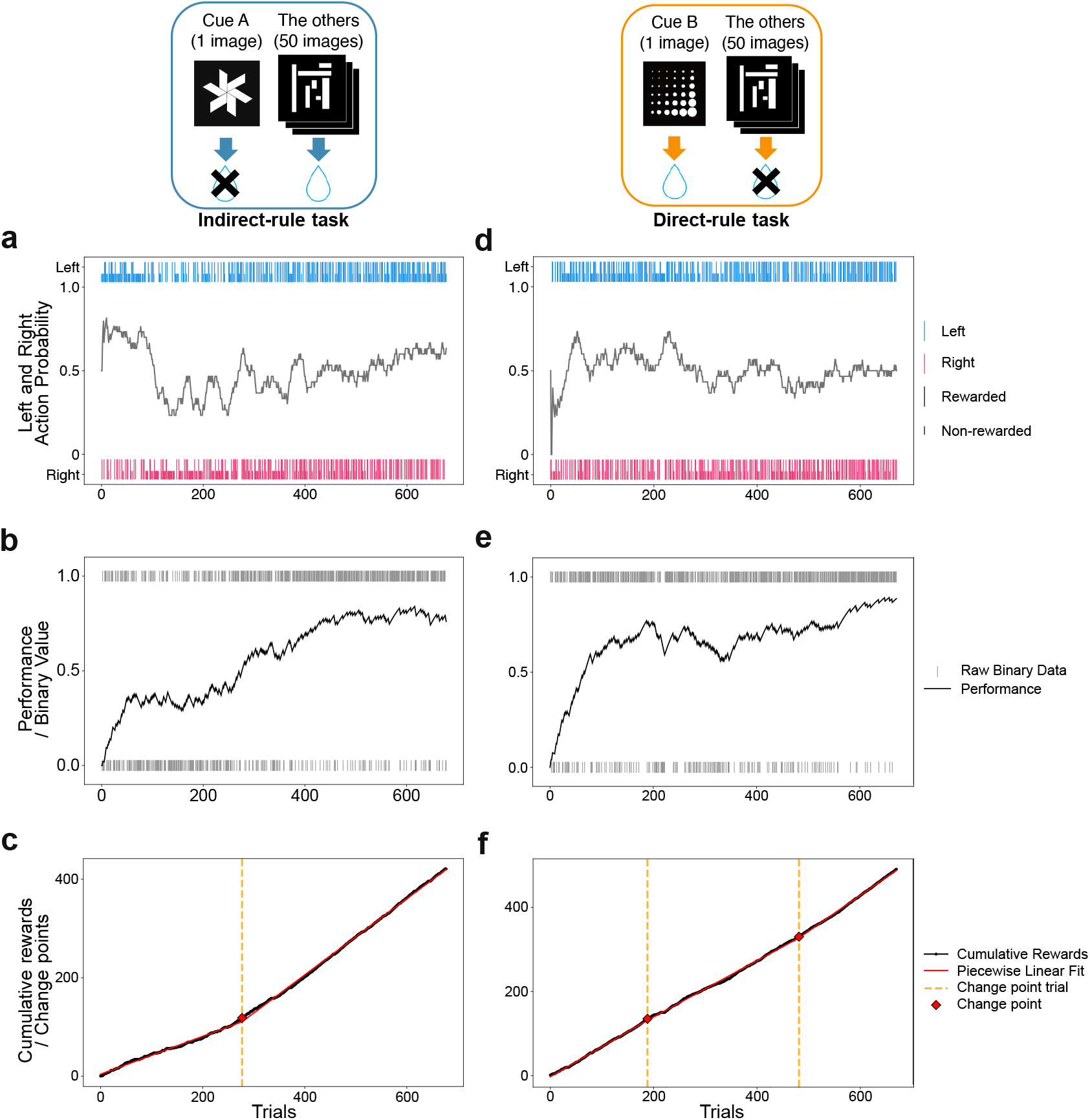
Behavioral transitions in the indirect- and direct-rule tasks. **a-c**, Representative behavioral session from a mouse performing the indirect-rule task. **d-f**, Representative behavioral session from a mouse performing the direct-rule task. **a**,**d**, Trial-by-trial left and right choice probabilities smoothed by an exponentially weighted moving average (EWMA). Blue and red lines indicate the two choices, and gray tick marks indicate raw choices on individual trials. **b**,**e**, Trial-by-trial correct/incorrect outcomes and their EWMA-smoothed accuracy. Gray tick marks indicate raw binary outcomes on individual trials. **c**,**f**, Cumulative rewards across trials. A piecewise linear fit was applied to each cumulative reward trajectory to estimate the behavioral breakpoint at which the reward accumulation rate changed. The breakpoint is indicated by the orange dashed line and red diamond.

These behavioral changes suggest that learning involved a change in how animals interpreted the task structure. We interpreted the early phase as behavior that may be described by stronger reliance on a side-dependent world model, whereas the later phase may be described by stronger reliance on a cue-dependent world model. We therefore used these overt behavioral transitions as a starting point for modeling the latent dynamics of world-model reliance over trials.

### Decision-making with competing world models

To account for abrupt behavioral transitions during learning, we hypothesized that the animals’ choices were generated by a dynamic interplay between competing internal world models rather than by a single fixed model. Based on this idea, we constructed an agent decision-making model in which action selection arises from the weighted integration of two candidate world models: a side-dependent model and a cue-dependent model. In the side-dependent model, reward is assumed to be generated from latent states corresponding to side-dependent reward probabilities. Under this model, the agent attempts to infer the reward probabilities associated with the left and right choices. In the cue-dependent model, reward is assumed to be generated from latent states corresponding to cue-dependent reward probabilities. Under this model, the agent attempts to infer the reward probabilities associated with the specific cue and with the other cues (**Fig. 1b, left**).

In each world model, the observed reward is generated from an underlying latent variable *z*, with reward probability given by a sigmoid function, 1/(1 + *e*^−*z*^). The agent was assumed to regard this latent variable as a hidden state that evolves according to random-walk dynamics over time, i.e., *z*_*t*_ = *z*_*t* −1_ + noise. Based on such a world model, the agent sequentially updated its belief about the latent reward structure in a Bayesian manner, following a free-energy-based formulation developed in our previous work [13] (**see Methods**). Specifically, the posterior belief over the latent state at trial t was approximated by a Gaussian distribution,

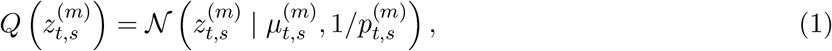

where *m* ∈ {side, cue}denotes the world model. In the side-dependent model, *s* ∈ {left, right}, whereas in the cue-dependent model, *s* ∈ {specific, other}.

In addition to these latent states, we introduced an internal latent variable *w*_*t*_ representing the degree to which the agent relied on each world model (**Fig. 1b left and bottom**). Under this formulation, the agent relied on the side-dependent model with weight 1 −*σ*(*w*_*t*_) and on the cue-dependent model with weight *σ*(*w*_*t*_), where *σ*(·) denotes the sigmoid function. At each trial, the agent was assumed to consider both world models and to choose the option with the higher expected reward probability based on their weighted integration. Thus, action selection was modeled as a mixture of decisions derived from the two competing world models.

The agent was assumed to update these beliefs sequentially in a Bayesian manner. The posterior mean of the latent state followed the update rule

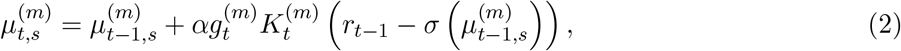

where *α* is the learning rate, 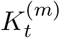 is the Kalman-gain-like coefficient, *r*_*t*−1_ is the reward outcome at the previous trial, and *σ*(·) maps the latent state to reward probability.

Here, 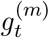 represents a gating term that modulates learning according to the current reliance on each world model (**Fig. 1c**). The rationale for this assumption is that a world model should not be strongly updated when the agent does not currently rely on it. In the gated learning model, 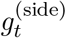 was set to 1 −*σ*(*w*_*t*_) and 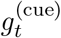 was set to *σ*(*w*_*t*_), so that updating of a world model was attenuated when the agent assigned little reliance to that model. Details of the component-specific belief updates, trial timing, and gating formulation are described in Methods.

### Inside insight dynamics method

Here, we developed a machine-learning method, termed inside insight dynamics (IID), to infer latent internal dynamics underlying insight-like behavioral transitions from observable action and reward sequences (**Fig. 1c**). IID introduces an inverse state-space model from the observer’s perspective, which we refer to as the observer-SSM, to infer the hidden internal states of the agent that give rise to observable behavioral sequences.

The observer-SSM assumes that the animal’s behavior is generated by the internal Bayesian decision-making model described above, in which two competing world models, a side-dependent model and a cue-dependent model, jointly govern decision-making. These models are characterized by latent reward states, 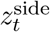 and 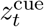, and by an internal weighting variable *w*_*t*_ that controls the relative influence of the two world models on behavior. Based on the observed action–reward history, the observer-SSM infers, at each trial, the hidden variables from the observer’s perspective: namely the estimated reward probabilities under each world model, the confidence associated with those estimates, and the relative reliance on the competing world models. The inference was implemented using a particle filter (see Methods). Through this framework, IID enables the observer to track latent restructuring of internal world-model dynamics during learning.

To validate the ability of IID to recover latent internal dynamics, we next performed simulation-based analyses. We first specified a latent insight variable *w*_*t*_ and used the forward model to generate behavioral sequences. In this simulation, actions were generated trial by trial, and the agent’s beliefs about reward probability within each world model were updated sequentially according to the resulting action–reward history. We then applied IID to the simulated behavioral data alone, without observing the ground-truth trajectories of *w*_*t*_ or the underlying belief states. The inferred latent dynamics were in good agreement with the ground truth, confirming that IID can recover both the insight-related weighting dynamics and the associated belief-updating process from behavior alone (**Supplementary Fig. S4**).

### Inverse estimation of insight dynamics in indirect-rule task

Applying IID to mouse behavioral data from the indirect-rule task, we estimated latent insight dynamics, including the mouse’s reward-probability estimates under each world model 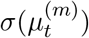, the confidence associated with those estimates 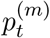, and the time-varying reliance on the competing world models (**Fig. 3**).

**Fig. 3.**
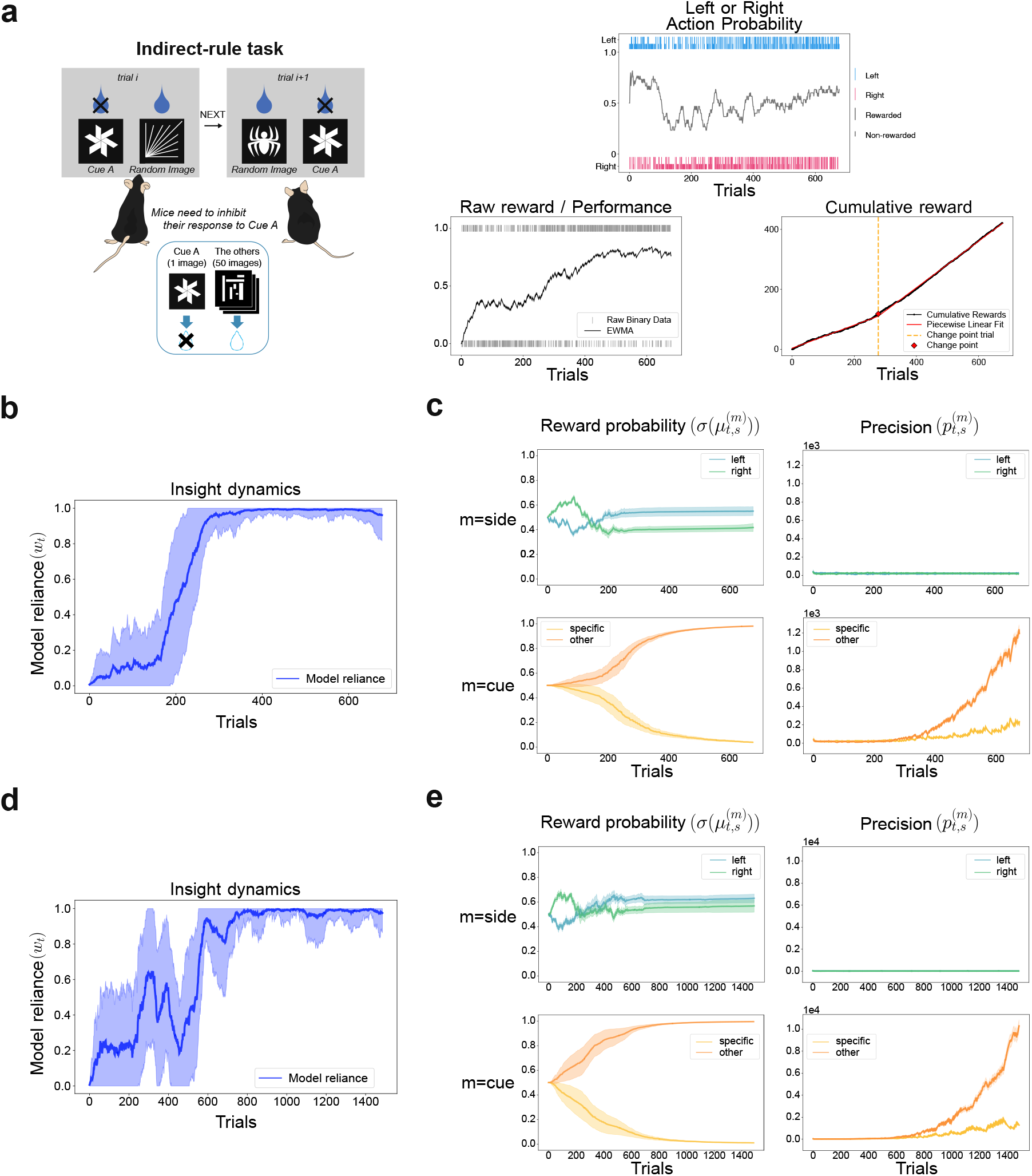
Inverse estimation of insight dynamics in a representative indirect-rule task sessions. **a**, Schematic of the indirect-rule task structure shown in **Fig. 1a** and behavioral data from a representative mouse performing the indirect-rule task shown in the same format as **Fig. 2a-c**. From left to right, the panels show the indirect-rule task schematic, EWMA-smoothed left/right choice probabilities with raw choices, EWMA-smoothed accuracy with raw binary outcomes, and cumulative reward with piecewise linear fit. The behavioral breakpoint estimated from the cumulative reward trajectory is indicated by the orange dashed line and red marker. **b**, Inferred insight dynamics for the mouse shown in **a**. The plot shows the estimated latent weighting variable *σ*(*w*_*t*_), representing the trial-by-trial reliance on the cue-dependent world model relative to the side-dependent world model. The orange dashed line indicates the behavioral breakpoint estimated from the cumulative reward trajectory in a. **c**, Inferred belief/recognition states within each world model for the mouse shown in **a**. Left panels show estimated reward probabilities, and right panels show the corresponding precisions. Upper and lower rows correspond to the side-dependent and cue-dependent world models, respectively. These inferred trajectories show that the increase in *w*_*t*_ was accompanied by the formation of cue-dependent reward beliefs and increased confidence in those beliefs during learning. **d, e**, Inferred insight dynamics and belief/recognition states in a different mouse performing the indirect-rule task from the mouse shown in **a–c**. The same format as **b** and **c** is used.

The estimated latent states indicated that this mouse underwent an abrupt transition at around trial 200, characterized by a rapid shift in the relative reliance on the competing world models (**Fig. 3b**). This latent transition was accompanied by an abrupt increase in the slope of the cumulative reward curve, consistent with a sudden improvement in reward acquisition (**the rightmost panel of Fig. 3a**). Notably, the transition detected by IID preceded the change point identified from the cumulative reward dynamics, suggesting that IID captures latent restructuring of the internal world model before it is fully manifested in overt behavior.

We next examined how latent reward beliefs evolved within each world model (**Fig. 3c**). Early in learning, updating was dominated by the side-dependent model, reflecting the animal’s stronger reliance on this model at the beginning of the task. Under the side-dependent model, the estimated reward probabilities for the left and right choices fluctuated around 0.5, consistent with the fact that reward was obtained with approximately equal probability on the two sides. Confidence in these estimates remained modest throughout this phase, indicating that the side-dependent model did not provide a highly reliable account of the task structure.

Around trial 100, however, the animal began to assign non-negligible weight to the cue-dependent model, suggesting that it had started to consider the possibility that reward might depend on cue identity rather than choice side. Because learning was gated by the current reliance on each world model, this shift allowed the cue-dependent model to be updated more strongly. Accordingly, the estimated reward probabilities under the cue-dependent model became increasingly differentiated. Around trial 200, when reliance on the two models became comparable (*w*_*t*_ ≈ 0.5), the cue-dependent model rapidly converged on the correct task structure: the reward probability associated with the specific cue approached 0, whereas that associated with the other cues approached 1.0. At the same time, confidence in the cue-dependent estimates increased sharply, indicating the formation of a reliable internal representation of the task structure.

As another example of IID estimation, we examined a second representative mouse performing the indirect-rule task (**Fig. 3d, e**). In this mouse, the estimated reliance on the cue-dependent model did not exhibit a clear abrupt transition, but instead increased gradually with a temporary decrease before eventually stabilizing near the cue-dependent state (**Fig. 3d**). The corresponding belief trajectories nevertheless showed increasing separation of cue-dependent reward probabilities, together with a later increase in precision (**Fig. 3e**). These results suggest that IID can capture not only clear insight-like transitions, but also more gradual and non-monotonic changes in world-model reliance, in which competition between alternative models persists before convergence.

### Gated learning better explains insight dynamics in indirect-rule task

Our analyses thus far have assumed a gated learning mechanism, whereby learning in each world model is weighted by the animal’s current reliance on that model. Under this view, once the animal begins to assign partial weight to a new world model, learning within that model proceeds in proportion to its internal weight. This formulation captures the idea that animals selectively update hypotheses that are currently considered relevant, rather than updating all possible interpretations of the task equally.

However, an alternative possibility is that multiple world models are learned in parallel, irrespective of their current contribution to behavior. This parallel learning mechanism can be implemented as a non-gating account (*g*_*t*_ = 1), in which the animal updates both candidate world models simultaneously, even before one of them is expressed strongly enough to guide action selection. Distinguishing between gated learning and parallel learning is important because it addresses whether insight-like transitions arise from reliance-dependent updating of an emerging world model, or from the later behavioral expression of a world model that has already been learned in parallel (**Fig. 4a**).

**Fig. 4.**
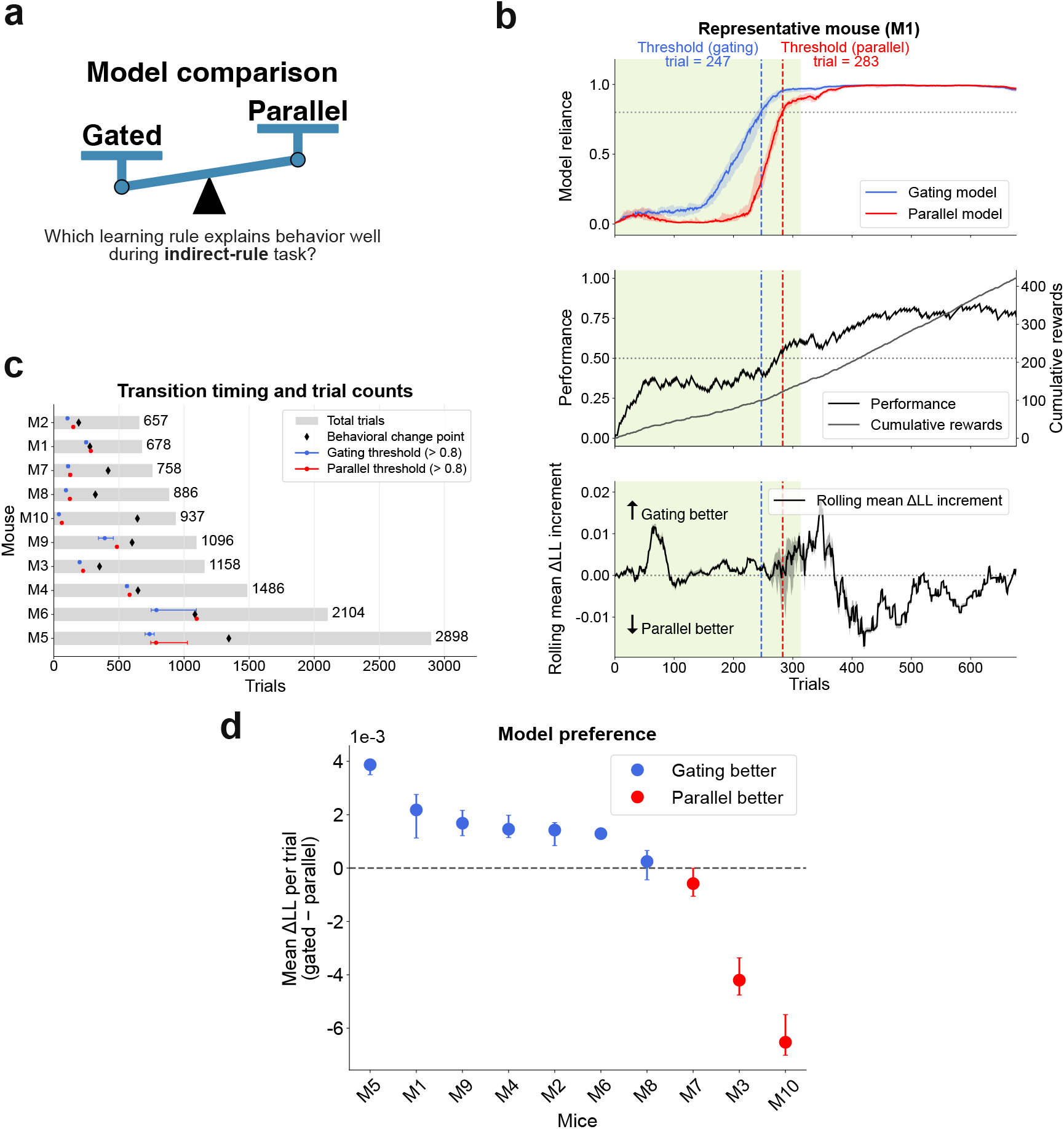
Model comparison between gated and parallel learning in the indirect-rule task. **a**, Schematic of the model comparison. We compared two alternative learning mechanisms: a gated learning model, in which belief updating in each world model is weighted by the current reliance on that model, and a parallel learning model, in which the two world models are updated in parallel. **b**, Representative comparison between gated and parallel learning models in one mouse. This representative mouse corresponds to the indirect-rule task example shown in **Fig. 2a-c** and **Fig. 3**, and is labeled M1 in c and d. The upper panel shows inferred insight dynamics under the gated learning model and the parallel learning model. Dashed vertical lines indicate the inferred insight timing in each model, defined as the trial at which the inferred insight variable exceeded 0.8. The middle panel shows behavioral performance and cumulative rewards. The lower panel shows the moving average of the trial-wise predictive log-likelihood difference between the gated and parallel learning models. Shaded regions indicate variability across repeated inference runs with different random seeds. **c**, Insight timing across mice. Gray bars indicate the total number of trials for each mouse. Black diamonds indicate change points in the slope of the cumulative reward trajectory detected by piecewise linear fitting. Blue and purple markers indicate the inferred insight timing under the gated and parallel learning models, respectively. Insight timing was defined as the trial at which the inferred insight variable exceeded 0.8. **d**, Summary of model preference across mice. Y-axis shows the mean trial-wise predictive log-likelihood difference between the gated and parallel learning models, defined as gated minus parallel. Positive and negative values indicate better action-prediction performance by the gated and parallel learning models, respectively. Points indicate medians across repeated inference runs with different random seeds, and asymmetric error bars indicate the 5th and 95th percentiles across seeds.

To resolve this issue, we compared the gated learning model and the parallel learning model based on their predictive performance for observed action sequences in the indirect-rule task. For each model, we inferred insight dynamics and computed the trial-wise predictive log-likelihood of action selection. In a representative mouse, both models captured an increase in the insight variable, but their inferred insight timing and action-prediction performance differed (upper panel in **Fig. 4b**).

As a behavioral reference, we also detected the change point in the slope of the cumulative reward trajectory by piecewise linear fitting, together with the corresponding behavioral performance trajectory (middle panel in **Fig. 4b**). We then quantified model preference over time using the trial-wise predictive log-likelihood difference between the two models (lower panel in **Fig. 4b**).

Across mice, we compared the inferred insight timing from the two models with this cumulative-reward change point (**Fig. 4c**). To summarize model preference for each mouse, we averaged the predictive log-likelihood difference over time. The gated learning model showed better action-prediction performance in 7 of 10 mice (**Fig. 4d**). Individual mouse-level model-comparison trajectories for the indirect-rule task, including seed-dependent variability and the transition-defined analysis epochs, are shown in (**Supplementary Fig. S5**). As a complementary visualization of model evidence over time, cumulative trial-wise log-likelihood advantages for the indirect-rule task are shown in **Supplementary Fig. S6**. These results support the view that, in the indirect-rule task, learning of the cue-dependent world model is better explained by a gated learning mechanism than by parallel learning of both candidate models. This interpretation is consistent with the structure of the indirect-rule task, in which successful performance requires discovery of a hidden and indirect cue-based rule rather than straight-forward reinforcement of an already explicit cue–reward association.

### Direct-rule task behavior favors parallel learning over gated learning

We next applied IID to the direct-rule task, where successful performance can be achieved by learning a more direct cue–reward relationship than in indirect-rule task (**Fig. 5a**). This allowed us to ask whether gated learning was still required when the cue-based rule was more directly accessible. We compared the gated and parallel learning models using the same likelihood-based procedure as in the indirect-rule task.

**Fig. 5.**
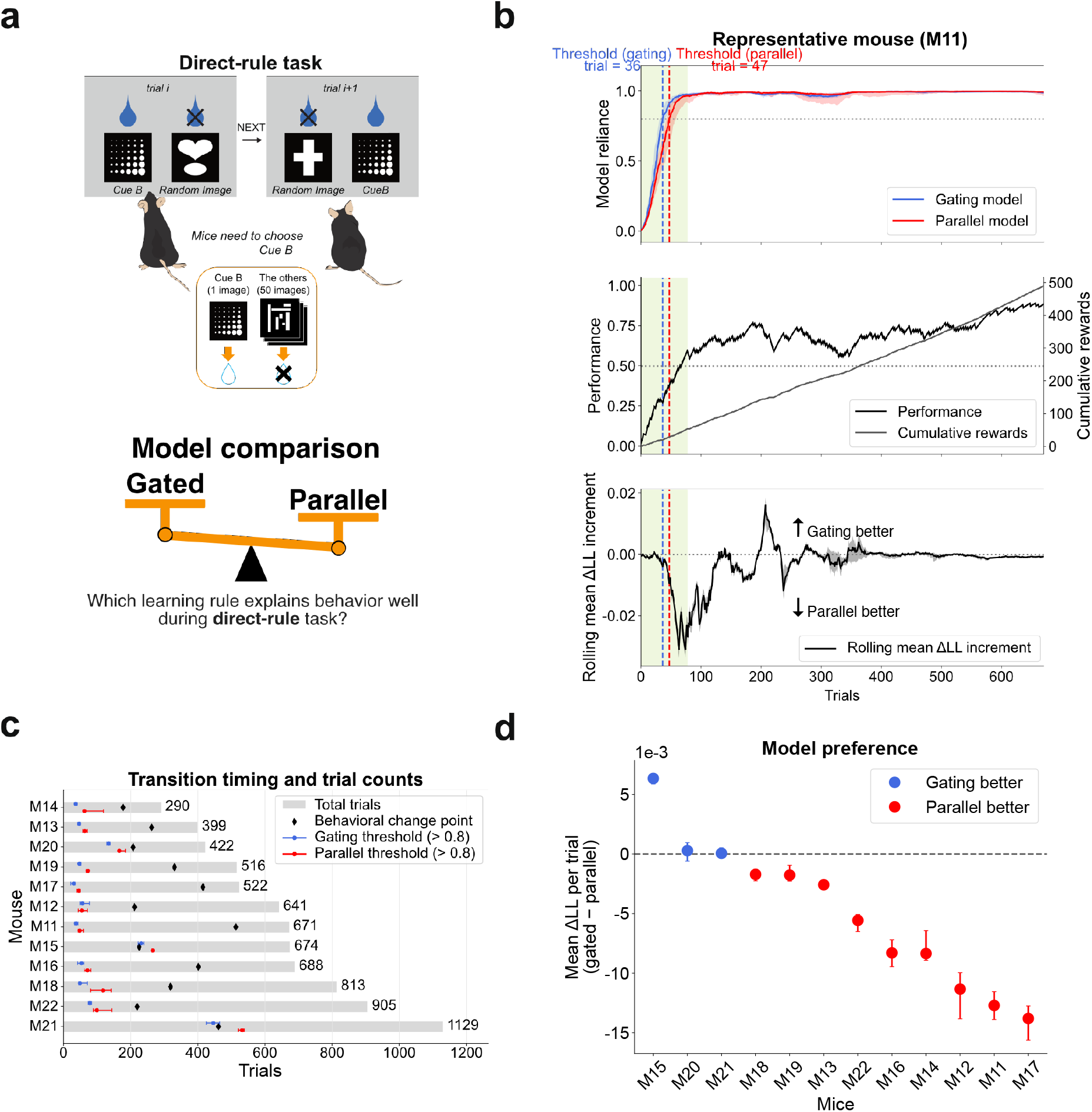
Model comparison between gated and parallel learning in the direct-rule task. **a**, Schematic of the direct-rule task structure shown in **Fig. 1a** and comparison between gated and parallel learning models, as in **Fig. 4a. b**, Representative comparison in one mouse performing the direct-rule task. This representative mouse corresponds to the direct-rule task example shown in **Fig. 2d-f**, and is labeled M11 in **b-d**,. The upper panel shows inferred insight dynamics under the gated and parallel learning models, with dashed lines indicating inferred insight timing, defined as the trial at which the insight variable exceeded 0.8. The middle panel shows behavioral performance and cumulative rewards, and the lower panel shows the moving average of the trial-wise predictive log-likelihood difference between the two models. Shaded regions indicate variability across repeated inference runs with different random seeds. **c**, Inferred insight timing across mice. Gray bars indicate total trial counts, black diamonds indicate change points in the slope of cumulative reward trajectories, and blue and purple markers indicate inferred insight timing under the gated and parallel learning models, respectively. **d**, Summary of model preference across mice. Y-axis shows the mean trial-wise predictive log-likelihood difference between the gated and parallel learning models, defined as gated minus parallel. Positive and negative values indicate better action-prediction performance by the gated and parallel learning models, respectively. Points indicate medians across repeated inference runs with different random seeds, and asymmetric error bars indicate the 5th and 95th percentiles across seeds.

In a representative mouse, both models inferred a transition toward the cue-dependent world model during learning, although the inferred insight trajectories differed between the gated and parallel learning models (upper panel in **Fig. 5b**). To relate these latent dynamics to overt behavior, we also plotted behavioral performance and cumulative rewards for the same session (middle panel in **Fig. 5b**). We then quantified model preference over time using the trial-wise predictive log-likelihood difference between the two models (lower panel in **Fig. 5b**).

Across mice, we compared the inferred insight timing from the two models with the cumulative-reward change point (**Fig. 5c**). The inferred insight points, as well as the cumulative-reward change point, appeared earlier in the direct-rule task than in the indirect-rule task (compare **Fig. 5c with Fig. 4c**). This suggests that mice reached the cue-based solution earlier in direct-rule task, regardless of how the transition timing was quantified.

We next asked which learning mechanism better explained these earlier transitions in direct-rule task (**Fig. 5d**). The gated learning model did not show a consistent predictive advantage. Instead, the parallel learning model achieved a higher action-prediction performance in 9 of 12 mice. Individual mouse-level model-comparison trajectories for the direct-rule task are shown in (**Supplementary Fig. S7**). Cumulative trial-wise log-likelihood advantages for the direct-rule task are shown in **Supplementary Fig. S8**. Thus, direct-rule task behavior was not consistently better explained by gated learning, suggesting that the more direct cue–reward structure may allow the cue-dependent world model to be acquired through parallel learning.

Together with the indirect-rule task results, these findings reveal a task-dependent dissociation in the computational organization of insight dynamics. In indirect-rule task, insight-like transitions were more consistently explained by gated learning, whereas in direct-rule task this gating advantage was not observed. This contrast indicates that the process by which animals acquire a new world model is not fixed, but depends on the structural demands of the task. Importantly, IID made it possible to uncover this distinction from behavioral data by reconstructing latent internal dynamics that would otherwise remain inaccessible.

## Discussion

In this study, we developed inside insight dynamics (IID), a computational framework based on the premise that an agent may have multiple competing world models and that learning depends on how these models are explored, updated, and selected. Within this framework, we operationally defined insight-like change as a transition between internal world models. We applied IID to mouse two-choice tasks in which successful performance required a shift between competing world models, and found that the resulting abrupt behavioral transitions are better understood not simply as changes in performance, but as latent restructuring of the internal world models through which animals interpret task structure. In this sense, the present work provides a quantitative approach for studying insight-like behavioral change as an internal restructuring process rather than as an externally observed shift in performance alone.

A central question in this study is what computational mechanism underlies the inferred restructuring of internal world models. In particular, the IID method allowed us to distinguish between two alternatives: a gating mechanism, in which learning within each world model depends on the animal’s current reliance on that model, and a parallel learning mechanism, in which multiple candidate world models are updated in parallel. Our results showed that the answer depended on task structure. In the indirect-rule task, behavior was better explained by the gating formulation, suggesting that a newly emerging world model became learnable only as the animal began to assign weight to it. By contrast, in the direct-rule task, the parallel learning formulation provided a better account, consistent with the idea that the more direct cue–reward structure of this task allows competing world models to be updated in parallel. Thus, the present study does not merely identify latent restructuring as a substrate of insight-like behavioral change, but further shows that the computational mechanism of that restructuring is itself task dependent.

To clarify what is meant by “insight” in the present study, it is important to distinguish the present framework from the subjective “Aha!” experience often discussed in human psychology [2, 3]. This distinction is especially important in animal cognition, where insight-like behavior cannot be validated through the phenomenology that defines the human “Aha!” experience and where the relation between human insight and putative nonhuman insight remains conceptually disputed [14, 15]. Human studies of insight have often relied on subjective report to identify the moment of insight, but such reports may not fully capture the temporal structure of the underlying process [16, 17]. In particular, under a gating mechanism, a newly emerging world model becomes learnable gradually as the agent begins to assign weight to it, potentially before the moment at which the subject consciously reports having reached a solution. IID may therefore provide a way to quantify a latent computational process that precedes subjective report or contributes to it, potentially extending the study of insight beyond subjective report alone.

Although the present study did not directly analyze neural activity, IID is particularly important for future neuroscience studies using animal models, such as mice, rats, and monkeys, in which insight-like change cannot be accessed through subjective report. Neural data are often interpreted in relation to observable behavior, but behavioral readouts alone may be insufficient for understanding what neural activity represents unless the latent internal dynamics underlying behavior are also taken into account [6, 18]. In the present case, these dynamics correspond to insight-like restructuring, including changes in the relative weighting of competing internal world models and the confidence associated with them. Relating such latent variables to neural activity may make it possible to identify neural processes that mediate or reflect the emergence of insight-like change.

A key methodological feature of the present study is that it treats behavior as arising from time-varying internal states rather than from fixed latent parameters [19, 20]. Computational modeling has provided powerful tools for inferring hidden cognitive variables from behavioral data, from trial-by-trial models of decision-making to Bayesian and active-inference frameworks for learning under uncertainty [8, 21–23]. Within this broader context, the distinctive aim of IID is not simply to estimate latent variables within a predefined task model, but to infer how the internal model space itself changes over time. This inverse perspective builds on our previous efforts to infer hidden computational variables from observable behavior. Earlier studies estimated relatively fixed latent quantities, such as reward-based behavioral strategies from animal behavioral time-series data [24] and participant-specific trust-related decision-making parameters from human trust-game behavior [25]. We later extended this inverse approach to time-varying internal states, including curiosity or internally weighted information seeking in two-choice behavior [13], and optimism-pessimism bias in monkey risk-taking behavior [26]. The present study shares this inverse perspective, but differs in one crucial respect: these previous studies assumed a single world model, whereas here we explicitly assume multiple competing world models and infer transitions between them. Thus, the main conceptual advance of IID is not only that it infers time-varying internal states, but that it decodes restructuring at the level of the internal model space itself. This makes IID particularly suited to studying abrupt representational change.

A major limitation of the present study is that the framework assumes behavior to arise from competition between only two candidate world models. This simplification was useful for the tasks studied here and allowed the latent restructuring process to be inferred in an interpretable manner, but the true internal hypothesis space of the animal may be richer than the binary alternatives considered here [6, 27]. Related to this, the specific forms of the candidate world models were also assumed rather than exhaustively derived from the data. For example, in the indirect-rule task, we considered a side-dependent model and a cue-dependent model, but other possible internal hypotheses may also have contributed to behavior. In particular, one could imagine alternative models in which reward is associated trial by trial with the randomly presented cue, rather than with the abstract category of “specific cue versus others.” Such possibilities were not explicitly modeled here, in part to maintain interpretability and computational tractability. The present results should therefore be understood as conditional on the candidate model set considered here.

In summary, this study introduces IID as a framework for inferring latent insight dynamics from behavior and shows that abrupt behavioral transitions can be understood as internal restructuring of competing world models. Our results further reveal that the computational form of this restructuring depends on task structure, with gating learning mechanisms dominating in a indirect-rule task and parallel learning mechanisms dominating in a more direct-rule task. By making latent restructuring quantitatively inferable from behavior alone, IID opens a new avenue for studying a central yet previously inaccessible aspect of flexible cognition. We hope that this framework will help establish a more quantitative understanding of insight-like change and, ultimately, provide a foundation for linking behavior, internal computation, and brain dynamics in the study of adaptive intelligence.

## Methods

### Behavioral tasks

Behavioral data analyzed in this study were derived from trial-level data underlying the previously published mouse visual discrimination study by Nishioka et al.[12]. The data were provided by the original investigators through collaboration for the purpose of the present secondary analysis. No new behavioral experiments were performed in this study.

The analyzed dataset consisted of male C57BL/6J mice that performed either the VD-Inhibit task or the VD-Attend task during behavioral learning. Mice were 8-10 weeks old at the beginning of the original behavioral experiments. The present analysis included 10 mice in the VD-Inhibit condition and 12 mice in the VD-Attend condition. In the dataset analyzed here, VD-Inhibit and VD-Attend were performed by separate cohorts of mice, and no animal contributed data to both tasks. The present study focused exclusively on trial-by-trial sequences of choices and reward outcomes during behavioral acquisition and did not analyze neural recordings or intervention data reported in the original study.

Mice were tested in touchscreen-based two-choice visual discrimination tasks, termed VD-Attend and VD-Inhibit. The task procedures are briefly summarized here, with full experimental details reported in the original study [12]. In each trial, animals initiated the trial by nose-poking the reward magazine, after which two visual cues were presented on the touchscreen until a response was made. A touch response to one of the two response windows determined the trial outcome. Correct responses were rewarded, whereas incorrect responses resulted in a timeout penalty. Training sessions consisted of up to 60 trials or 60 min per day, and mice were trained until they reached the criterion of more than 80% correct responses for two consecutive days. In the VD-Attend task, a fixed visual cue (the marble image) served as the rewarded target, whereas responses to the alternative image, randomly selected from a predefined image set, were unrewarded and punished by a timeout. In the VD-Inhibit task, by contrast, responses to a fixed visual cue (the flag image) were designated as incorrect, whereas responses to the alternative image were rewarded. Thus, the VD-Attend task involved learning a direct cue-reward association, whereas the VD-Inhibit task required animals to suppress responses to a fixed non-rewarded cue and instead select the alternative image.

### Behavioral data and preprocessing

Behavioral data were analyzed on a trial-by-trial basis for each mouse. For each trial, we extracted the animal’s choice and the corresponding binary reward outcome. Raw image identities were not used as model inputs. Instead, cue-related information was reduced to the task-relevant category, specific cue versus other cue, according to the fixed task rule and the recorded choice–outcome relation. Because each trial contained one fixed cue and one alternative cue, this mapping allowed the chosen and unchosen options to be assigned to the corresponding cue-dependent components used in the model. Specifically, we tracked whether the chosen option corresponded to the task-relevant target or non-target category, as determined by the reward contingency of each task. Rewarded trials were coded as 1 and unrewarded trials as 0. These trial-wise action and reward sequences were used both for descriptive behavioral analyses and as inputs to the inside insight dynamics (IID) method.

The analysis focused on the initial learning period of each task, from the first training session to the session in which the animal reached the learning criterion. Sessions after criterion acquisition were not included in the main IID analysis. Trials with missing action or reward records were excluded from analysis. Animals were included if trial-level choice and reward-outcome records were available throughout the learning period required for IID inference. No additional animals were excluded after applying these criteria.

For visualization purposes, reward sequences were smoothed by an exponentially weighted moving average,

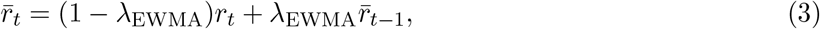

where *r*_*t*_ denotes the binary trial outcome and *λ*_EWMA_ is reported in Supplementary Table S1. Choice probabilities were computed using a causal sliding 30-trial window,

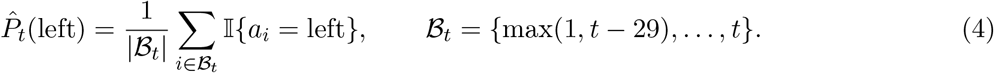

The corresponding right-choice probability was defined as 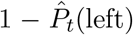. These smoothing procedures were used only for visualization and descriptive behavioral summaries. All IID analyses were performed on the original unsmoothed trial-by-trial action–reward sequences and the corresponding task-defined category mappings.

### Behavioral change-point analysis

As a conventional behavioral reference, we estimated behavioral change points from cumulative reward dynamics for comparison with the latent transition times inferred by IID. For each animal, cumulative reward was defined as the cumulative sum of binary reward outcomes across trials,

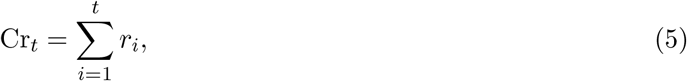

where *r*_*i*_ ∈{0, 1} denotes the reward outcome at trial *i*, and *t* = 1, …, *T*. We fitted the cumulative reward trajectory with a continuous two-segment piecewise-linear regression model, also known as a broken-line or segmented regression model [28]. Using trial number *t*, the fitted curve was written as

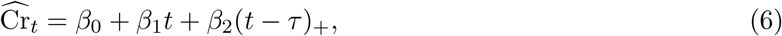

where (*u*)_+_ = max(*u*, 0), and *τ* denotes the internal breakpoint. This parameterization enforces continuity at the breakpoint, with the slope changing from *β*_1_ before *τ* to *β*_1_ + *β*_2_ after *τ*. The breakpoint and regression coefficients were estimated by minimizing the residual sum of squares,

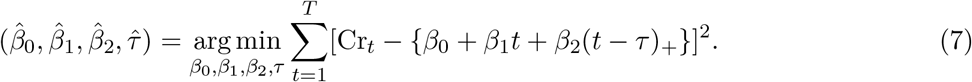

In the implementation, this fit was performed using the Python package `pwlf` with two line segments, corresponding to one internal breakpoint (version 2.0.0; https://github.com/cjekel/piecewise_linear_fit_py). The estimated internal breakpoint was taken as a descriptive behavioral change point, reflecting the trial at which the rate of reward acquisition changed in the cumulative reward trajectory. This behavioral change point was used only for comparison with IID-inferred transition times and not as an independent inferential test of a discrete latent state transition.

### Multi-world-model decision-making model

To model abrupt behavioral transitions, we assumed that the agent’s behavior was generated by competition between multiple internal world models. We considered two candidate models: a side-dependent model and a cue-dependent model. Insight-like change was defined as a transition in the relative dominance of these models.

### Agent-SSM for each world model

Let *m* ∈ {side, cue} denote the world model, and let *s* denote the component index within each model. For the side-dependent model, *s* ∈ {left, right}. For the cue-dependent model, *s* ∈ {specific, other}. For each world model, we assumed a latent reward state 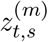 that evolved across trials according to a random-walk process,

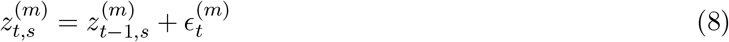

Where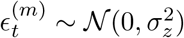 is Gaussian noise with variance 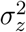.

Reward probability was expressed as

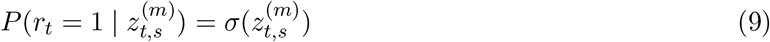

where *σ*(*x*) = 1/(1 + *e*^−*x*^), and *r*_*t*_ denotes the reward outcome at trial *t*.

### Approximate belief updating in the agent-SSM

Following our previous free-energy-based formulation, the agent’s belief about the latent reward state was approximated by a Gaussian distribution,

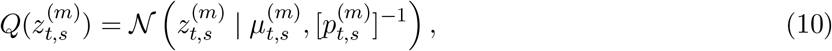

where 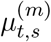 is the posterior mean and 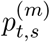 is the precision. Here, the latent variables at trial *t* denote the agent’s pre-choice belief state after incorporating the action and reward outcome from the preceding trial. Because reward feedback was obtained only for the chosen option, belief updating was component-specific. We introduced an indicator 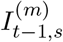 that equals 1 when component *s* of world model *m* was sampled on trial *t* − 1, and 0 otherwise. The posterior mean and precision were updated sequentially as

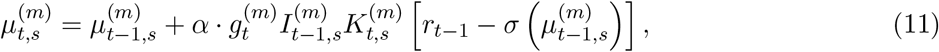

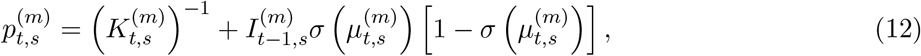

where *α* is the learning rate, 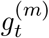 is the gating coefficient for the posterior-mean update by the current model reliance, and 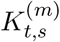 is the Kalman-gain-like coefficient defined as

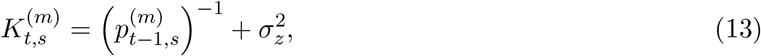

with 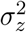 denoting the variance of the latent-state random walk. The gating term directly modulated only the reward-prediction-error update of the posterior mean. However, because the precision update for sampled components was evaluated using the updated posterior mean, gating can influence precision indirectly through its effect on 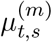. These updates followed the same free-energy-based approximation as in our previous work [13].

### Gated and parallel learning formulations

In the gating formulation, the posterior-mean update within each world model depended on the current reliance:

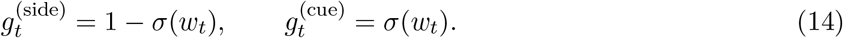

In the parallel learning formulation, the posterior means in both world models were updated independently of the current model reliance. This was implemented as a non-gated update rule:

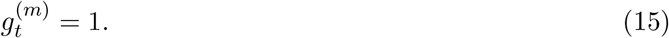

Thus, the key difference between the two formulations was whether reward-prediction-error updates of the posterior means were scaled by the agent’s current reliance on each world model. In the gated learning formulation, model reliance controlled the magnitude of the update, whereas in the parallel learning formulation, both world models were updated whenever their corresponding components were sampled, regardless of their current contribution to action selection.

### Action selection under competing world models

We introduced an internal weighting variable *w*_*t*_ to represent the relative dominance of the two world models. The side-dependent and cue-dependent models were weighted by 1 − *σ*(*w*_*t*_) and *σ*(*w*_*t*_), respectively.

For the side-dependent model, 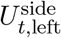 and 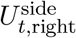 denote the agent’s estimated reward probabilities for choosing the left and right options, respectively. For the cue-dependent model, 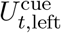 and 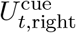 denote the estimated reward probabilities for the left and right options under the cue-dependent interpretation. Because the cue-dependent model is defined over *s*∈ {specific, other}, these quantities depend on which cue category is presented on each side at trial *t*. The integrated utilities for the left and right choices were defined as

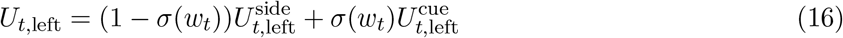

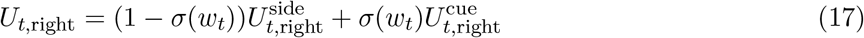

Here,

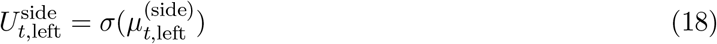

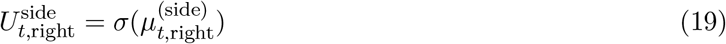

and, depending on cue placement,

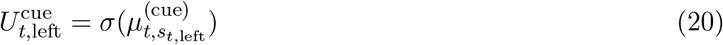

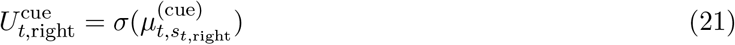

where *s*_*t*,left_, *s*_*t*,right_ ∈ {specific, other}. Assuming *a*_*t*_ ∈ {0, 1} where 0 and 1 denote the left and right choices respectively, the action selection probability was then given by

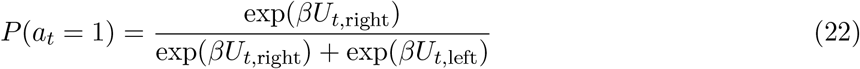

where *β* is the inverse-temperature parameter, which was consistently set to 1.0 in our model.

### Inside insight dynamics (IID) method

IID adopts the observer’s perspective to infer the hidden internal states of the agent under the multi-world-model decision-making framework described above.

### Observer-SSM

From the observer’s perspective, the latent state at trial *t* was defined as

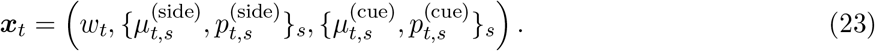

where *w*_*t*_ denotes the relative weighting of the competing world models, and 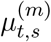 and 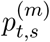 denote the posterior mean and precision of the latent reward belief in world model *m*, which are updated according to (Eqs. (11) and (12)), respectively. Here, *x*_*t*_ denotes the agent’s pre-choice internal state at trial *t*, after incorporating the action and reward outcome from trial *t*− 1. Thus, the current action *a*_*t*_ was generated from *x*_*t*_, whereas the reward outcome *r*_*t*_ was used to update the belief state for the next trial.

The hidden-state dynamics were assumed to follow

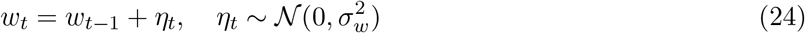

where 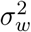 denotes the variance of the system noise. The belief variables 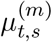and 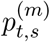 evolved according to the agent-side belief updating equations described above. Thus, the observer-SSM combined stochastic dynamics of the world-model weighting variable with deterministic, component-specific updates of the latent reward beliefs. Additional details on trial timing and component-specific belief updating are provided in **Supplementary Note 1**. The observable variables at trial *t* were the agent’s action *a*_*t*_ and reward outcome *r*_*t*_. The action likelihood was given by Eq. (22), while the reward outcome entered the state transition from trial *t* to trial *t* + 1.

### Sequential inference by particle filtering

Using the observer-SSM, the posterior distribution of the agent’s latent internal state ***x***_*t*_ was estimated sequentially from the observed behavioral data (*a*_*t*_, *r*_*t*_). The latent state at trial *t* was inferred in a Bayesian manner as

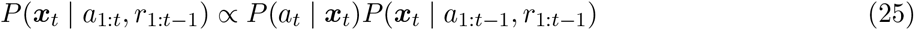

where *a*_1:*t*_ denotes the action history up to trial *t*, and *r*_1:*t*_−_1_ denotes the reward history up to the preceding trial. The predictive distribution was given by

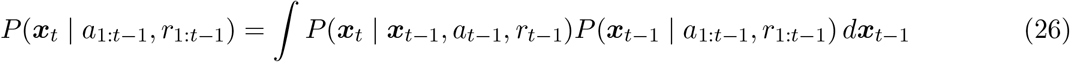

where *P* (***x***_*t*_| ***x***_*t*−1_) is the latent-state transition model defined by the observer-SSM. The predictive prior incorporated the state transition induced by the previous trial’s action and reward, whereas the current action likelihood provided the evidence for the latent state at the time of choice.

Because this posterior cannot be obtained analytically owing to the nonlinear and non-Gaussian structure of the model, we approximated it using a particle filter[29–31]. Specifically, the posterior distribution was represented by a set of weighted particles,

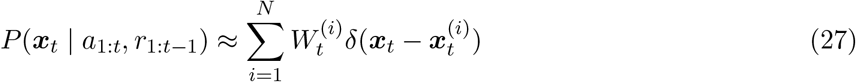

where 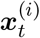 is the *i*-th particle, 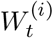is its normalized importance weight, *N* is the number of particles, and *δ*(·) denotes the Dirac delta function. Within each particle, the components of the latent state were updated according to the observer-SSM, including the random-walk dynamics of the world-model weighting variable *w*_*t*_ (Eq. (24)) and the belief-updating equations for 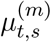 and 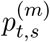 (Eqs. (11) and (12)). At each trial, particles were propagated according to the latent-state dynamics and reweighted by the likelihood of the observed action. Because resampling was performed adaptively, the normalized weights carried by the particle system immediately before the update at trial *t* were denoted by 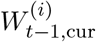. If resampling had been performed after the previous update, then 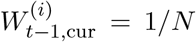 for all particles; otherwise, 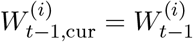

For the bootstrap particle filter used here, particles were sampled from the latent-state transition model,

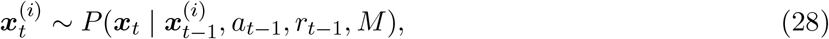

so that the incremental log weight was given by the current action likelihood,

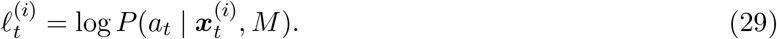

The unnormalized importance weight was therefore computed as

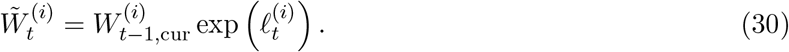

Equivalently, in log space,

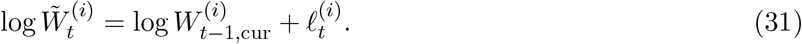

The incremental predictive log-likelihood at trial *t* was computed from the normalizing constant of this weight update,

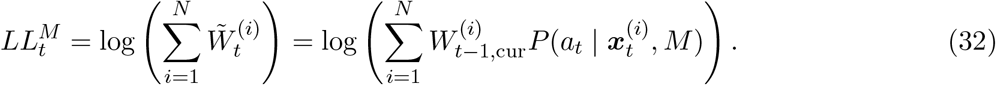

For numerical stability, this quantity was evaluated in log space as

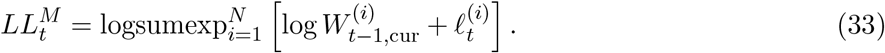

The accumulated predictive evidence of the observed action sequence was tracked separately as the sum of the incremental predictive log-likelihoods,

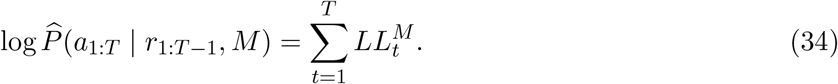

The normalized importance weights were then updated as

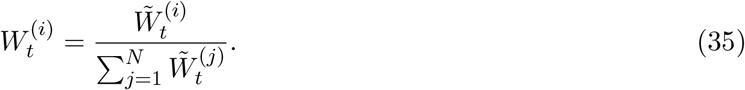

Particle degeneracy was monitored using the effective sample size[32, 33],

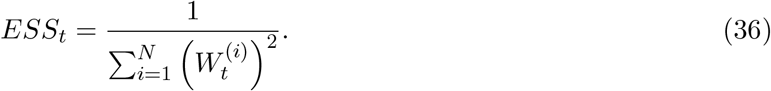

Adaptive resampling was performed when *ESS*_*t*_/*N* fell below the prespecified threshold. After resampling, all particles were assigned equal normalized weights, 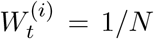, and their ancestral indices were stored. If the threshold was not crossed, resampling was skipped and the nonuniform normalized weights were carried forward to the next trial. Thus, when resampling was skipped, the previous normalized weights contributed explicitly to the next trial’s unnormalized weights; when resampling was performed, this contribution was already represented by the ancestor-selection probabilities.

To summarize path-space latent trajectories, we reconstructed ancestral paths for terminal particles by tracing the stored ancestral indices backward over the full trial sequence. For each terminal particle *i*, we computed an additive observation-fit score

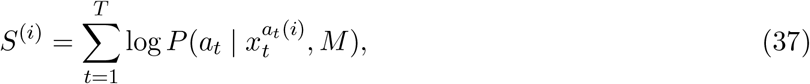

where *a*_*t*_(*i*) denotes the particle index at trial *t* along the ancestral lineage of terminal particle *i*. This score can be viewed as an additive path functional of the latent trajectory, a class of quantities widely studied in particle smoothing and online smoothing methods.[34, 35] Representative paths were then selected from a prespecified empirical quantile band of this score. The displayed latent dynamics were summarized across the retained ancestral paths using normalized within-band selection weights,

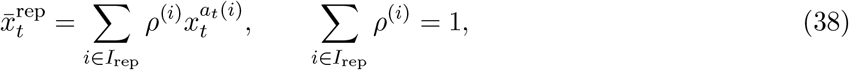

where *I*_rep_ denotes the set of retained terminal-particle indices and *ρ*^(*i*)^ denotes the normalized aggregation weight assigned to retained path *i*. In the implementation used for visualization, path stratification was based on the cumulative observation-fit score, and the displayed trajectories were given by weighted within-band expectations of the latent states and their sigmoidal transforms.

This path-selection procedure was used to obtain representative latent trajectories for visualization and for defining transition-defined analysis epochs. The trial-wise predictive log-likelihood was estimated from the full weighted particle system produced by the particle filter rather than from the representative trajectories. In the model-comparison analysis, the threshold-crossing times used to define the analysis epoch were computed from the representative model-reliance trajectories obtained by this path-selection procedure. Accordingly, path selection contributed to the model-comparison analysis through the definition of the temporal window over which predictive log-likelihood differences were summarized.

### Simulation-based validation of IID

To validate the ability of IID to recover latent internal dynamics, we performed simulation-based analyses using the forward model described above. Specifically, we first prescribed trial-by-trial trajectories of the latent weighting variable *w*_*t*_, and then generated behavioral sequences from the multi-world-model decision-making model. Under this process, actions were generated trial by trial, while the latent beliefs within each world model, represented by 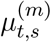 and 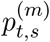, were sequentially updated according to the resulting action–reward history. We then applied IID to the simulated behavioral data alone, without access to the ground-truth trajectories of *w*_*t*_, 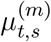, or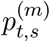. This procedure allowed us to test whether the observer-SSM could recover the latent internal dynamics that had generated the behavior.

### Model comparison between gated and parallel learning models

Model comparison consisted of two related but distinct steps. First, for each animal, model, and random seed, we estimated the trial-wise predictive log-likelihood of the observed actions using the full particle-filtering procedure described above. Second, to summarize model preference during early learning and the immediate post-transition period, we averaged the trial-wise predictive log-likelihood difference within a transition-defined analysis epoch. This epoch was defined using a threshold-crossing rule applied to the representative model-reliance trajectories obtained by the path-selection procedure described above.

Let 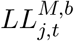 denote the trial-wise predictive log-likelihood for animal *j*, model *M* ∈ gated, parallel, and random seed *b*. The trial-wise predictive advantage of the gated model over the parallel model was defined as

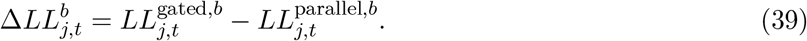

For each animal, model, and random seed, the representative model-reliance trajectory was obtained from the retained ancestral paths selected by the cumulative observation-fit score. The inferred transition time was then defined as the first trial at which this representative reliance on the cue-dependent world model exceeded 0.8. Let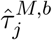 denote this threshold-crossing trial. In all analyzed runs, the inferred reliance exceeded the threshold within the session, so a threshold-crossing trial was obtained for every seed and model.

To define a single analysis epoch for each animal, we first summarized the seed-dependent transition times within each model by their median:

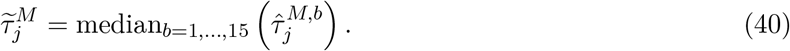

The analysis epoch was then defined from the beginning of the session to 30 trials after the later of the median transition times under the gated and parallel learning models:

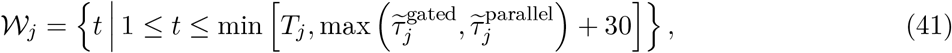

where *T*_*j*_ is the total number of analyzed trials for animal *j*. We used the later of the two model-specific transition times to ensure that the analysis epoch covered the inferred transition period under both candidate models.

For each seed, model preference was quantified as the average predictive log-likelihood difference within this window:

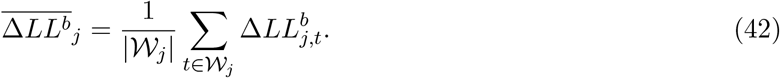

Positive and negative values indicate better predictive performance of the gated and parallel learning models, respectively. For visualization and summary statistics, the point estimate for each animal was defined as the median of 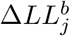 across seeds, and across-seed variability was summarized by the 5th and 95th percentiles.

Because the per-animal model-preference summary was evaluated within an analysis epoch defined from path-selected model-reliance trajectories, it should be interpreted as a transition-defined predictive-performance comparison rather than as an epoch-independent comparison over the entire behavioral sequence.

### Analysis parameters and implementation settings

All numerical parameters used for behavioral preprocessing, IID inference, particle filtering, simulation-based validation, and model comparison are summarized in Supplementary Table S1. Unless otherwise stated, the same parameter settings were used for the VD-Inhibit and VD-Attend analyses. Parameters that were fixed rather than fitted were chosen before the final model comparison and were kept identical across animals and across the gated and parallel learning formulations.

## Data availability

The behavioral data analyzed in this study were derived from the trial-level data of the previously published study by Nishioka et al. [12]. The processed trial-by-trial datasets used for IID analysis, including action and reward-outcome sequences for each animal, will be made publicly available at [repository/DOI] upon publication.

## Code availability

The code used for behavioral preprocessing, IID inference, simulation-based validation, model comparison, and figure generation was implemented in Python (v3.12). The analysis code, parameter files, and instructions required to reproduce the main figures and supplementary analyses will be made publicly available at [repository/DOI] upon publication.

## Ethics statement

No new animal experiments were performed in this study. This study involved secondary analysis of mouse behavioral data that had been collected for, and used in, a previously published study by [12]. The original animal experiments were conducted in accordance with the guidelines of the National Institutes of Health and were approved by the Animal Experimental Committee of the Institute for Protein Research, The University of Osaka (approval IDs: 29-02-1 and R04-01-0). The present study reanalyzed these existing data provided by the original investigators through collaboration, and no additional animal procedures were conducted.

## Acknowledgements

This work was supported in part by the Japan Science and Technology Agency (JST) Moonshot R&D Program (JPMJMS2024-9 to H.N.), JST CREST (JPMJCR25Q2 to H.N.), JST BOOST (JPMJBS2424 to K.I.), Japan Agency for Medical Research and Development (AMED) Multidisciplinary Frontier Brain and Neuroscience Discoveries (Brain/MINDS 2.0) (JP25wm0625322 to T.H. and JP25wm0625210 to H.N.), AMED Grant Numbers JP21wm0425010 and JP21gm1510006 (to T.H.), MEXT/JSPS KAK-ENHI Grant Numbers JP21K15209 (to T.N.), JP21K15210 (to T.M.), JP16H06568, JP18H02542, JP21H05694, JP22H02944, JP22H00494, and JP25K02547 (to T.H.), JSPS, Overseas Research Fellowships (to T.N.), Takeda Life Science Research Foundation (to T.H.), and the Collaborative Research Program of the Institute for Protein Research, the University of Osaka, CR-26-03 (to T.H.).

## Author contributions

K.I., T.H., and H.N. conceived the project. K.I. and H.N. developed the computational framework.

K.I. performed the computational analyses with assistance from M.F. in data preparation. T.N. and T.M. conducted the mouse experiments and acquired the behavioral data. K.I. and H.N. wrote the manuscript and prepared the figures, with input from all authors. H.N. supervised the computational study. All authors reviewed and approved the manuscript.

