## Supplementary Information for "Inside insight: decoding how insight emerges from competing world models"

### Supplementary notes

#### Trial timing and component-specific belief updating in IID

Here we provide additional details on the trial timing and component-specific belief updating used in IID. The latent state $\boldsymbol{x}_t$ was defined as the agent's pre-choice internal state at trial $t$. Therefore, $\boldsymbol{x}_t$ already incorporates the action and reward outcome from the preceding trial, $(a_{t-1}, r_{t-1})$, but not the reward outcome from the current trial. The current action $a_t$ was generated from this pre-choice state, and the reward outcome $r_t$ was then used to form the next pre-choice state, $\boldsymbol{x}_{t+1}$.

For each world model $m$, only the component sampled by the animal's choice was updated on a given trial. In the side-dependent model, the sampled component was the chosen side. In the cue-dependent model, the sampled component was the cue category, either the specific cue or the other cue category, presented on the chosen side. We denote this component-specific sampling indicator by $I_{t,s}^{(m)}$:

$$I_{t,s}^{(\text{side})} = \mathbb{I}\{s = a_t\}, \tag{1}$$

for the side-dependent model, and

$$I_{t,s}^{(\text{cue})} = \mathbb{I}\{s = c_t(a_t)\}, \tag{2}$$

for the cue-dependent model, where $c_t(a_t) \in \{\text{specific}, \text{other}\}$ denotes the cue category presented on the side chosen at trial $t$.

The transition from $\boldsymbol{x}_t$ to $\boldsymbol{x}_{t+1}$ was then given by a reward-dependent update of the sampled component. The posterior mean and precision were updated according to the belief-updating equations (Eqs. (11) and (12)) described in the main text, with the trial indices shifted from $t$ to $t+1$ to explicitly represent the transition from the current pre-choice state to the next pre-choice state following observation of reward outcome $r_t$. Thus, the observation-dependent term was added only for the component sampled on that trial. For unsampled components, $I_{t,s}^{(m)} = 0$, and the precision changed only through state-transition dynamics specified by the latent-state update model.

In the gated learning formulation, the model-reliance modulated only the posterior-mean update:

$$g_t^{(\text{side})} = 1 - \sigma(w_t), \qquad g_t^{(\text{cue})} = \sigma(w_t). \tag{3}$$

In the parallel learning formulation, the posterior means in both world models were updated independently of the current model reliance:

$$g_t^{(m)} = 1. \tag{4}$$

Importantly, the precision update was not directly multiplied by the gating term in either formulation. This implementation reflects the assumption that model reliance controls the magnitude of the reward-prediction-error update in the posterior mean, whereas the component-specific precision update reflects whether the corresponding component was sampled on that trial. Thus, the distinction between gated and parallel learning was implemented at the level of the posterior-mean update, while precision dynamics can still differ between the two formulations through the updated mean-dependent term.

### Supplementary Tables

**Supplementary Table S1: Analysis parameters and implementation settings.** This table summarizes the dataset descriptors, preprocessing settings, agent-side decision-model parameters, observer-SSM initialization and dynamics, observer-SSM particle-filtering settings, and model-comparison settings used in the IID analyses. Gamma distributions are parameterized by shape $k$ and scale $\theta$.

| Category | Parameter | Symbol / name | Value used / initialization rule |
|---|---|---|---|
| Dataset | Number of VD-Inhibit mice | $n_{\text{Inhibit}}$ | 10 |
| Dataset | Number of VD-Attend mice | $n_{\text{Attend}}$ | 12 |
| Preprocessing | EWMA smoothing parameter | $\lambda_{\text{EWMA}}$ | 0.02 |
| Preprocessing | Choice-probability window | – | 30 trials |
| Agent decision model | World-model formulation | $M$ | Gating or non-gating |
| Agent decision model | Learning-rate parameter | $\alpha$ | Fixed at 0.1 |
| Agent decision model | Inverse temperature | $\beta$ | Fixed at 1.0 |
| Agent decision model | Latent reward-state noise standard deviation | $\sigma_z$ | Fixed at 0.3 corresponding to $\sigma_z^2 = 0.09$ |
| Observer-SSM | Initial reward-belief mean | $\mu_{0,s}^{(m)}$ | Initialized as 0.0 corresponding to $\sigma(\mu_{0,s}^{(m)}) = 0.5$ |
| Observer-SSM | Initial reward-belief precision | $p_{0,s}^{(m)}$ | $p_{0,s}^{(m,i)} \sim \text{Gamma}(k = 2.5, \theta = 1.0)$ |
| Observer-SSM | Initial model-weight variable | $w_0$ | $w_0^{(i)} \sim \mathcal{N}(-5, 0.1^2)$ |
| Observer-SSM | Model-weight noise variance | $\sigma_w^2$ | $(\sigma_w^2)^{(i)} \sim \text{Uniform}(0.2, 0.7)$ |
| Observer-SSM particle filter | Number of particles | $N$ | 1500 |
| Observer-SSM particle filter | Proposal distribution | – | Bootstrap proposal from observer-SSM transition |
| Observer-SSM particle filter | Observation likelihood | $P(a_t \mid \boldsymbol{x}_t, M)$ | Action likelihood from Eq.(22) |
| Observer-SSM particle filter | Resampling method | – | systematic resampling |
| Observer-SSM particle filter | ESS threshold | ESS/$N$ | 0.5 |
| Model comparison | Analysis epoch | $\mathcal{W}_j$ | From trial 1 to 30 trials after later median transition |
| Model comparison | Number of repeated inference runs | – | 15 seeds |
| Model comparison | Rolling-average window for $\Delta LL_t$ | – | 30 trials |
| Model comparison | Model-reliance threshold | $\sigma(w_t)$ | 0.8 |

### Supplementary figures

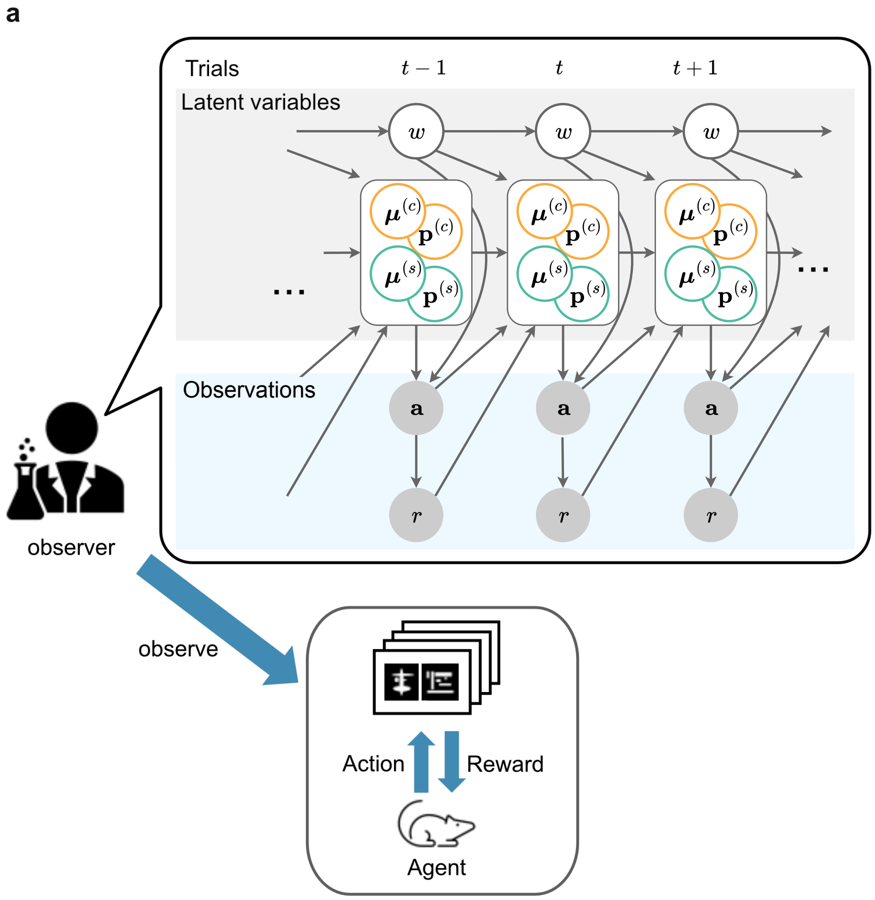


**Fig. S1 | Observer-SSM used in inside insight dynamics.**

The observer-SSM formulates the inverse problem of the agent decision-making model. The upper panel provides a graphical-model representation of the observer-side state-space model, showing how latent internal variables and behavioral observations are related across trials. The agent interacts with the task environment and generates trial-by-trial behavioral observations, consisting of actions $a_t$ and reward outcomes $r_t$. The observer receives only these observable sequences and infers the latent internal variables that most likely generated them. The latent state at each trial includes the world-model weighting variable $w_t$, representing the relative reliance on the cue-dependent versus side-dependent world model, and the belief states within each world model, represented by the posterior means $\mu^{(s)}, \mu^{(c)}$ and precisions $p^{(s)}, p^{(c)}$. In the gated formulation shown here, $w_t$ influences action selection and modulates the subsequent posterior-mean update within each world model. The precision update was component-specific but was not directly gated by $w_t$. By contrast, in the parallel learning formulation, the same action-selection process is used, but posterior-mean updating proceeds independently of $w_t$. Arrows indicate temporal dependencies among latent variables and the observation process linking latent internal states to actions and subsequent rewards. This observer-side formulation provides the basis for decoding insight dynamics from behavioral time-series data.

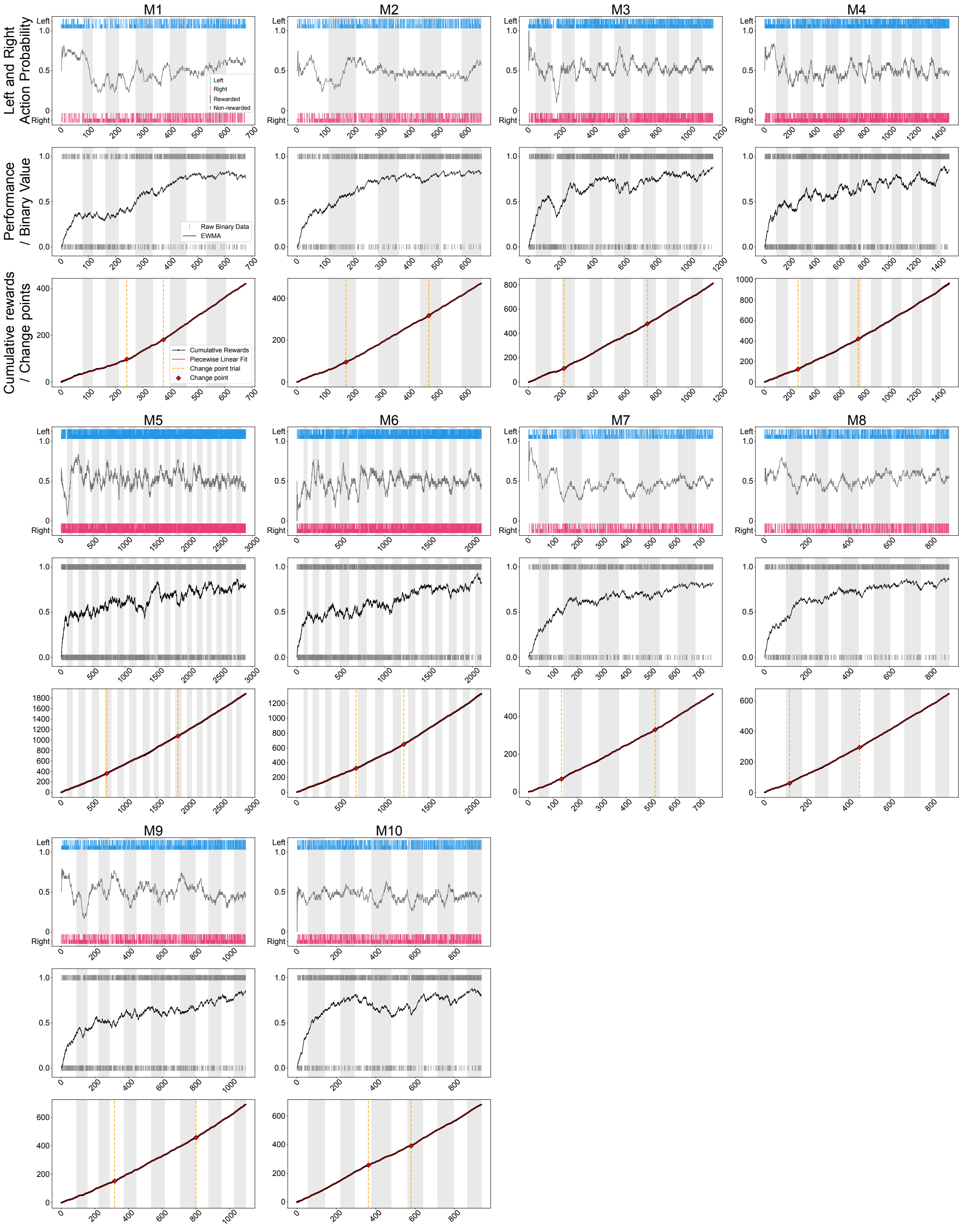


**Fig. S2 | Individual behavioral trajectories in VD-Inhibit mice.**

Individual behavioral trajectories for all VD-Inhibit mice, M1-M10. For each mouse, the upper panel shows trial-by-trial left and right choice probabilities, with raw choices indicated by tick marks. The middle panel shows binary trial outcomes and EWMA-smoothed behavioral performance. The lower panel shows cumulative rewards together with a two-segment piecewise-linear fit. Dashed vertical lines and red markers indicate behavioral change points estimated from the cumulative reward trajectory. Alternating white and gray background bands indicate consecutive session days along the concatenated trial axis; changes in background shading mark boundaries between session days.

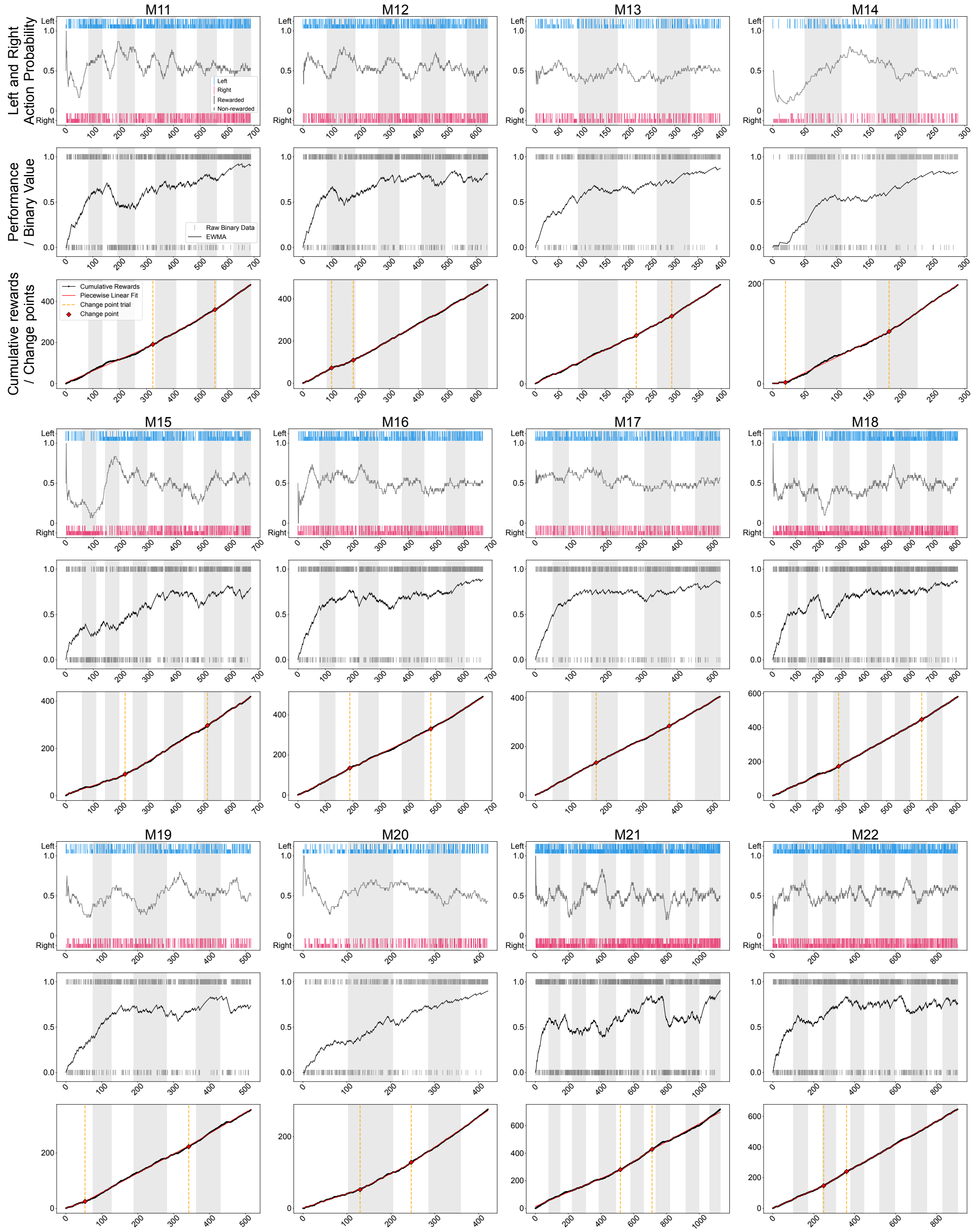


**Fig. S3 | Individual behavioral trajectories in VD-Attend mice.**

Individual behavioral trajectories for all VD-Attend mice, M11-M22. The format is the same as in **Fig. S2**, but applied to mice performing the VD-Attend task. For each mouse, the upper panel shows trial-by-trial left and right choice probabilities, the middle panel shows binary trial outcomes and EWMA-smoothed behavioral performance, and the lower panel shows cumulative rewards with a two-segment piecewise-linear fit. Dashed vertical lines and red markers indicate behavioral change points estimated from the cumulative reward trajectory. Alternating white and gray background bands indicate consecutive session days along the concatenated trial axis; changes in background shading mark boundaries between session days.

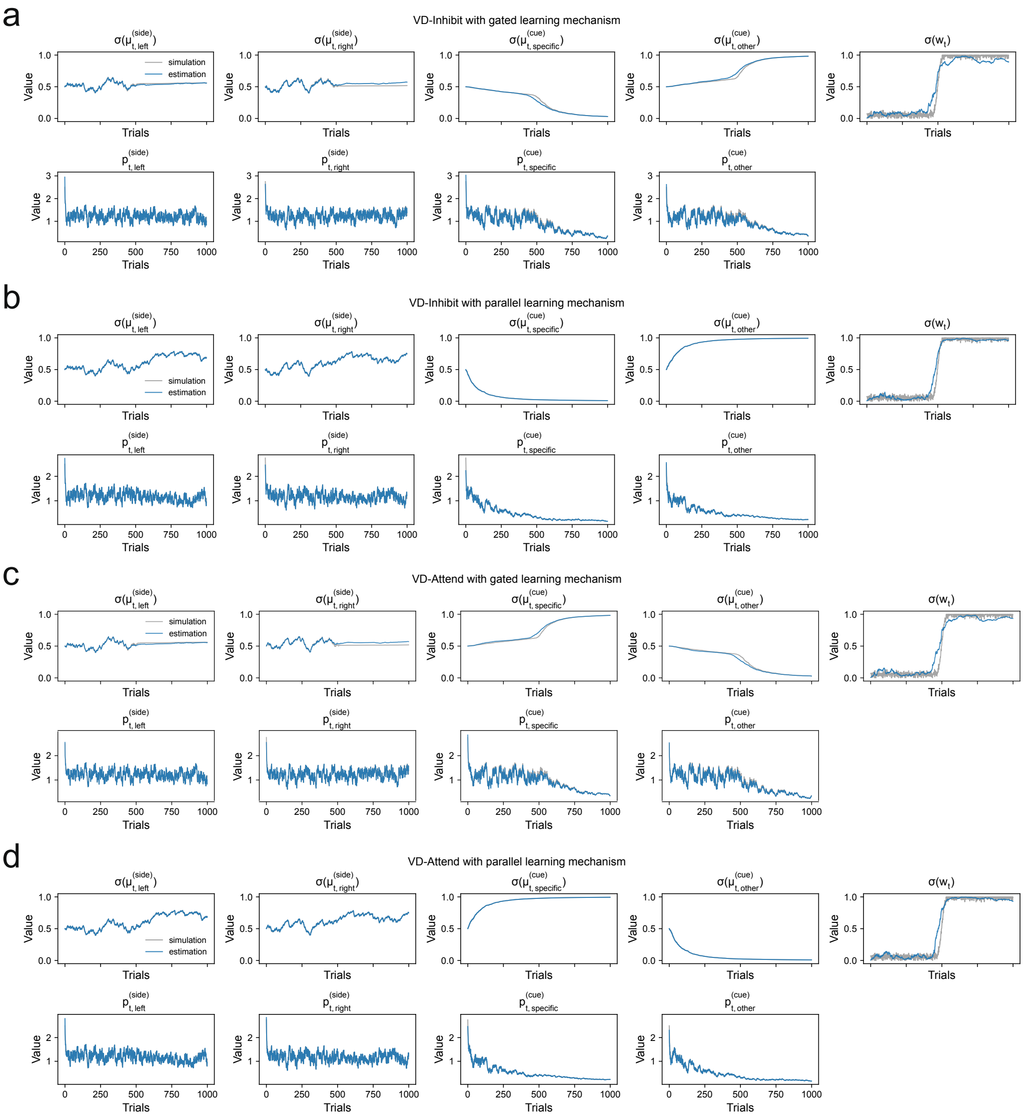


**Fig. S4 | Simulation-based validation of IID.**

**a-d,** Recovery of latent trajectories from synthetic action–reward sequences generated under four task and learning-mechanism conditions: VD-Inhibit with gated learning **a**, VD-Inhibit with parallel learning **b**, VD-Attend with gated learning **c**, and VD-Attend with parallel learning **d**. For each condition, synthetic behavior was generated from known latent trajectories and then analyzed with IID using only the generated choices, rewards, and task-defined category mappings. Gray lines indicate the simulated ground-truth trajectories, and blue lines indicate the trajectories estimated by IID. Top rows show sigmoid-transformed reward-belief variables for the side-dependent and cue-dependent world models, together with the sigmoid-transformed model-weight variable, $\sigma(w_t)$. Bottom rows show the corresponding precision variables for each belief dimension. Across task structures and learning mechanisms, IID recovered the overall temporal profiles of latent reward beliefs, precision dynamics, and model-weight transitions, supporting its ability to infer latent world-model dynamics from behavioral data.

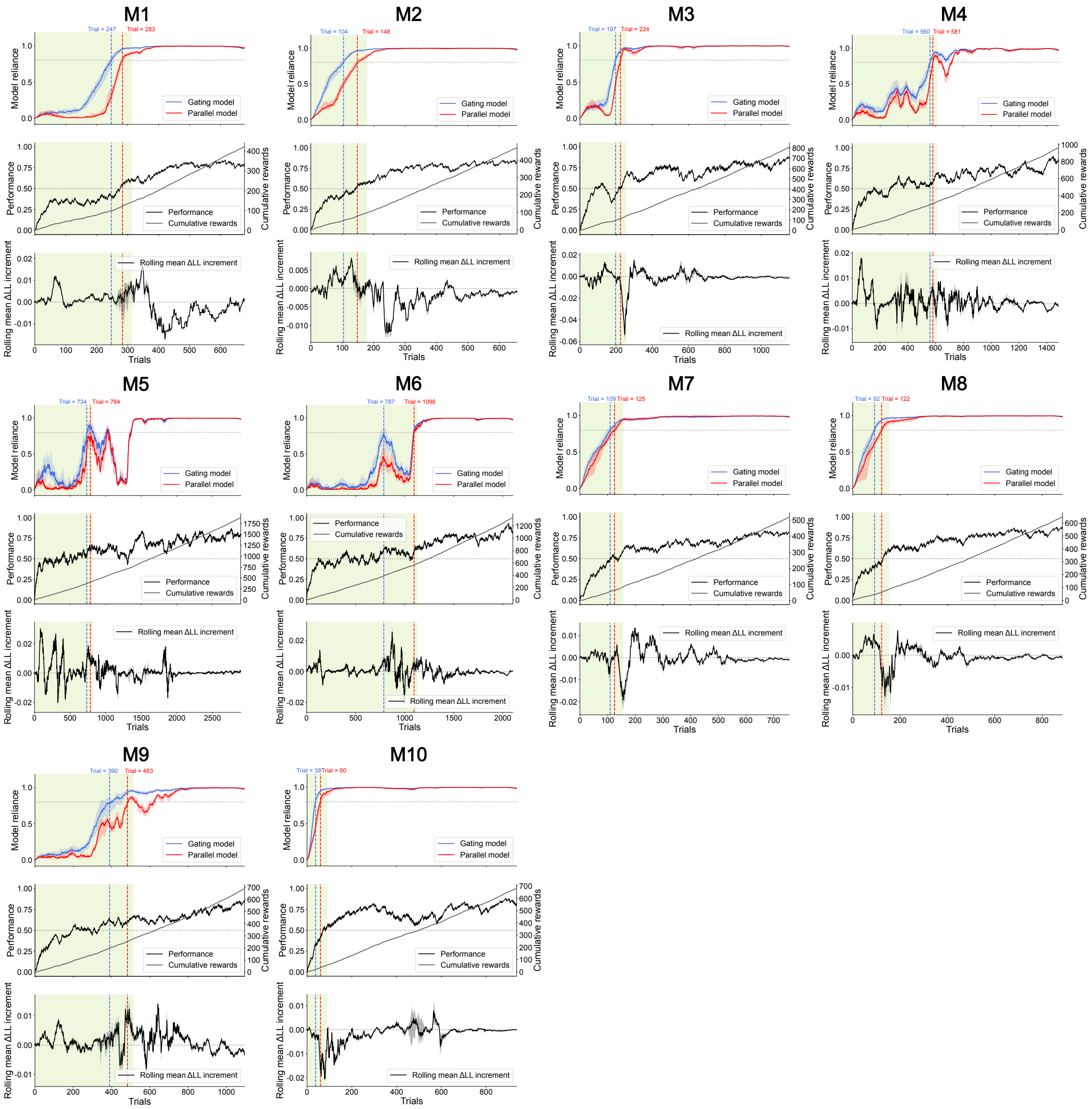


**Fig. S5 | Individual model-comparison results in VD-Inhibit mice.**

Individual results of the gated-versus-parallel model comparison for all VD-Inhibit mice, M1-M10. For each mouse, the upper panel shows estimated model reliance under the gated and parallel learning models. Colored bands surrounding the model-reliance trajectories indicate variability across repeated inference runs with different random seeds. Dashed vertical lines indicate the threshold-crossing trials, defined as the first trials at which the inferred reliance on the cue-dependent world model exceeded 0.8. The middle panel shows behavioral performance and cumulative rewards. The lower panel shows the rolling mean of the trial-wise predictive log-likelihood difference, $\Delta LL_t = LL_t^{\text{gating}} - LL_t^{\text{parallel}}$. Positive values indicate better predictive performance of the gated learning model, whereas negative values indicate better predictive performance of the parallel learning model. Pale green shaded regions indicate the analysis epochs used to summarize model preference for each mouse.

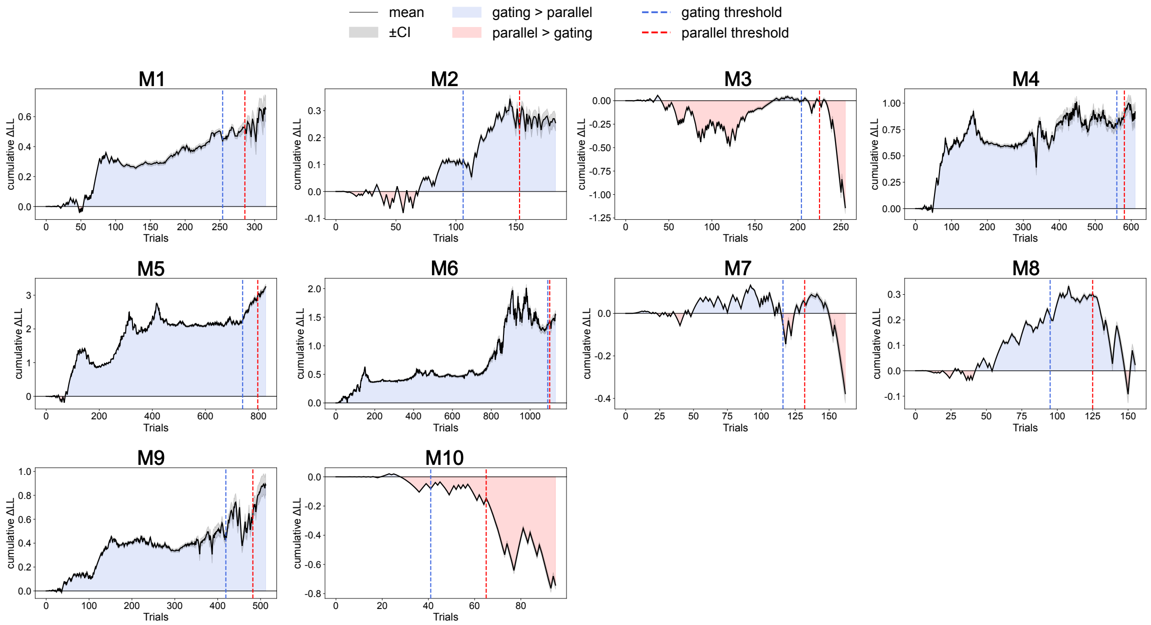


**Fig. S6 | Trial-wise cumulative log-likelihood advantage of gated versus parallel learning in the VD-Inhibit task.** For each mouse (M1-M10), the cumulative difference in observation log likelihood ($\Delta\mathrm{LL}$) between the gated-learning and parallel-learning models is plotted across trials. Axis ranges are scaled independently for each mouse to visualize within-mouse dynamics. Positive values indicate greater cumulative support for the gated-learning model, whereas negative values indicate greater cumulative support for the parallel-learning model. The black line denotes the across-seed mean, and the gray band denotes the corresponding confidence interval across particle-filter seeds. Blue and red background shading mark trial ranges in which the cumulative $\Delta\mathrm{LL}$ favors the gated-learning and parallel-learning models, respectively. Blue and red dashed vertical lines indicate the threshold-crossing times for the gated and parallel learning models. These threshold-crossing times were computed from representative trajectories obtained by the path-selection procedure and are overlaid here to relate model-comparison results to the analysis window, rather than being inferred from the cumulative $\Delta\mathrm{LL}$ curve itself.

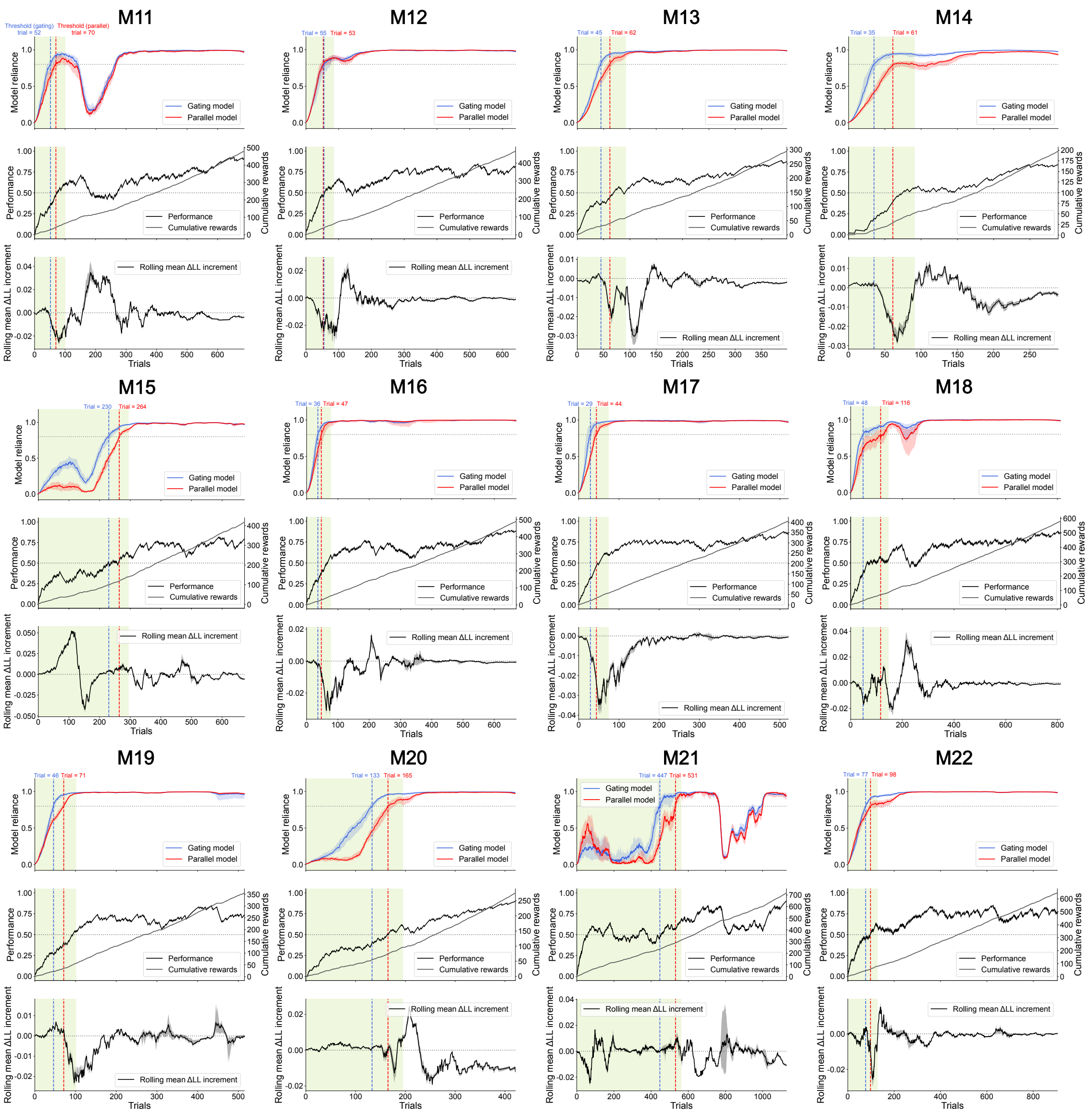


**Fig. S7 | Individual model-comparison results in VD-Attend mice.**

Individual results of the gated-versus-parallel model comparison for all VD-Attend mice, M11-M22. The format is the same as in **Fig. S5**, but applied to mice performing the VD-Attend task. For each mouse, the upper panel shows estimated model reliance under the gated and parallel learning models, with colored bands indicating variability across repeated inference runs with different random seeds and dashed vertical lines indicating threshold-crossing trials. The middle panel shows behavioral performance and cumulative rewards, and the lower panel shows the rolling mean of the trial-wise predictive log-likelihood difference, $\Delta LL_t = LL_t^{\text{gating}} - LL_t^{\text{parallel}}$. Pale green shaded regions indicate the analysis epochs used to summarize model preference for each mouse.

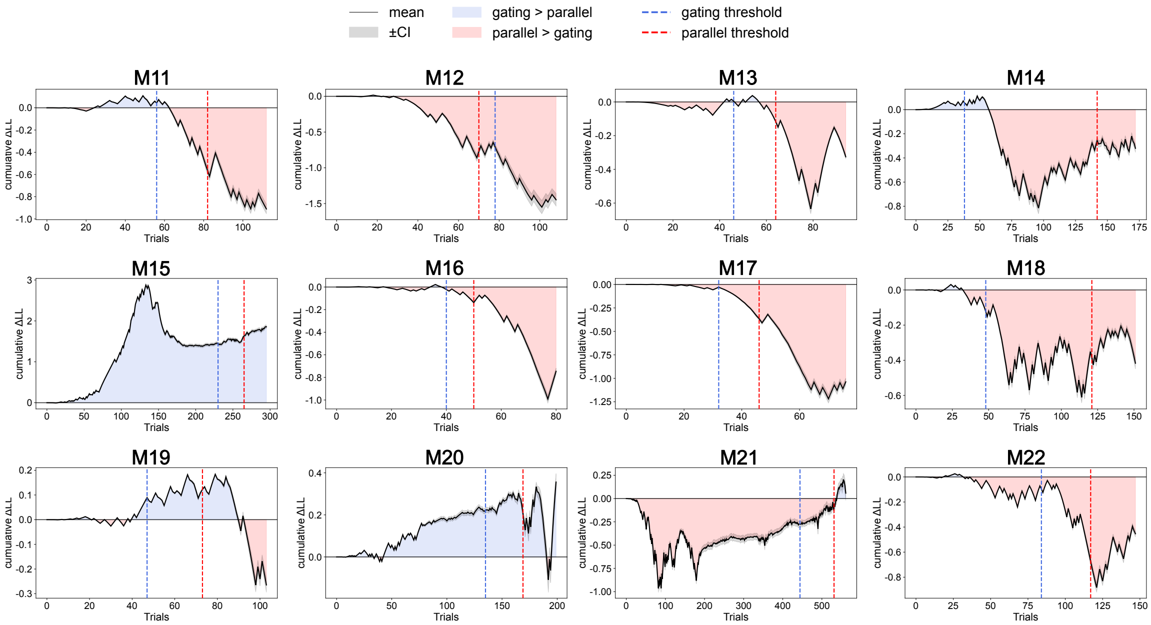


**Fig. S8 | Trial-wise cumulative log-likelihood advantage of gated versus parallel learning in the VD-Attend task.**
For each mouse (M11-M22), the cumulative difference in observation log likelihood ($\Delta\mathrm{LL}$) between the gated-learning and parallel-learning models is plotted across trials. Axis ranges are scaled independently for each mouse to visualize within-mouse dynamics. Positive values indicate greater cumulative support for the gated-learning model, whereas negative values indicate greater cumulative support for the parallel-learning model. The black line denotes the across-seed mean, and the gray band denotes the corresponding confidence interval across particle-filter seeds. Blue and red background shading mark trial ranges in which the cumulative $\Delta\mathrm{LL}$ favors the gated-learning and parallel-learning models, respectively. Blue and red dashed vertical lines indicate the threshold-crossing times for the gated and parallel learning models. These threshold-crossing times were computed from representative trajectories obtained by the path-selection procedure and are overlaid here to relate model-comparison results to the analysis window, rather than being inferred from the cumulative $\Delta\mathrm{LL}$ curve itself.
